# Characterizing antibodies binding to the same epitope reveals limited contribution by heavy chain CDR3 sequences relative to other CDRs

**DOI:** 10.1101/2025.05.24.655967

**Authors:** Jarjapu Mahita, Haeuk Kim, Marcus De Almeida Mendes, Eve Richardson, Jason A. Greenbaum, Alessandro Sette, Morten Nielsen, Bjoern Peters

## Abstract

The antigen specificity of antibodies is defined mainly by their highly-variable complementarity-determining regions (CDRs), of which the CDR3 is considered to be the most important. Similar antibodies are expected to bind the same epitope on an antigen, but how this similarity should be quantified is unclear. Here, we utilized a large body of publicly available data to identify features that are most informative in predicting if two antibodies are likely to bind to the same epitope. We examined features such as pairwise sequence identities, CDR structural similarity, and geometrical similarity of antibody paratopes. We found that, for antibodies with high overall sequence similarity, the CDR sequence identity alone is sufficient to predict if a pair of antibodies binds to the same epitope. Strikingly, the sequence identity of CDR1 and CDR2 had as much or more predictive power than CDR3, which is commonly thought of as the main determinant of antibody specificity. In addition, for antibody pairs with lower overall sequence identity, including structural information led to significant improvements in predictive performance. Guided by these results, we developed BCRMatch (https://github.com/IEDB/BCRMatch) which uses an ensemble machine-learning approach to predict if pairs of antibodies target the same epitope based on antibody sequence alone. This can be applied to identify potential targets of an antibody of interest by comparing it to antibodies with known specificities.

## INTRODUCTION

B cells are a major arm of the adaptive immune system, capable of generating highly diverse membrane-bound B-cell receptors (BCRs), which can be secreted as antibodies. A typical monomeric antibody is a Y-shaped protein complex containing two heavy and light-chain pairs. The high specificity of antibodies is mediated by their complementarity-determining regions (CDRs) which are also the main sources of variability in antibody sequences (*1*). Each heavy and light-chain encodes for three CDRs. The site of an antibody that binds to the antigen is known as the paratope and typically includes major portions of the CDRs. In contrast, the epitope is the region on the antigen that directly interacts with the antibody.

The diversity of BCRs arises due to two main processes. The first is somatic gene rearrangement of the variable (V), diversity (D) and joining (J) gene segments to form the heavy-chain V-domain (only V and J in the case of light-chain V-domain). The CDR1 and CDR2 are directly encoded in the V-gene segment while the CDR3 corresponds to the junction at which V(D)J (or VJ) gene segments are joined together, resulting in increased variability. The pairing of heavy-chain and light-chain at this stage results in a naïve BCR which has not yet been exposed to antigen.

Depending on the V, D, and J gene segments that comprise the V-domains of both chains, the naïve BCR can have pre-determined specificity (*2*). In the second phase, the naïve BCR binds an antigen, gets activated and subsequently migrates to the dark zone of the germinal center where it undergoes further mutation through a process called somatic hypermutation (SHM). This introduces random point mutations in the V-domains, mainly in the CDRs (*3*). B-cells having BCRs with mutations that increase affinity to the antigen are selected and may go through further rounds of SHM, eventually resulting in high-affinity antigen-specific antibodies (*4*).

Identifying the epitope specificity of antibodies is a widely researched topic. Traditional methods to experimentally characterize antibody epitopes entail isolating antigen-specific antibodies followed by binding assays and/or structural studies. The last decade has witnessed major advances made in high-throughput sequencing (HTS) technologies which has enabled their routine use for investigating antigen-specific immune responses (*5*-*15*). This has led to an exponential increase in available BCR/antibody sequences.

To mine antigen-specific and epitope-specific antibodies from this repertoire of sequences, several computational methods have been developed. These methods work mainly by clustering together antibodies with high sequence identity to one another (*7*), (*16* -*18*). High sequence similarity between antibodies can be caused by two factors: First, antibodies can originate from the same ancestral BCR and consequently be members of the same clonal lineage. Second, convergent evolution can lead antibodies from different clonal lineages to gain similarity – both in terms of sequence and structure - through somatic mutations, which is favored if they bind the same epitope (*19*-*22*).

In this work, we investigate which features of antibodies predict whether they bind to a shared epitope. We utilized sequence-and structure-based features to encompass all aspects relevant to antibody specificity and compared different machine-learning models using these features to predict shared epitope specificity. Based on these results, we present a newly developed tool ‘BCRMatch’, which accepts sequences of CDRs of antibodies and uses an ensemble of machine-learning models developed in this study to predict which antibodies share an epitope. Thus, our study provides quantitative insights into feature importance for antibody-epitope recognition and a directly applicable implementation of these findings into a freely available tool.

## RESULTS

### Assembling datasets of antibody pairs

To delineate the contributions of sequence and structural features of antibodies in predicting their specificity, we assembled three different datasets of antibody pairs along with their features. As detailed in the methods, we used three separate sources for obtaining variable heavy-chain (VH) and light-chain (VL) sequence and structural data on antibodies used to populate the datasets: 1) the Ab-Ligity study (*12*), 2) the Coronavirus Antibody Database (Cov-AbDb) (*13*), and 3) the Immune Epitope Database (IEDB, www.iedb.org) (*14*,*15*). The Ab-Ligity study also has a pre-constructed dataset of antibody pairs and their corresponding class labels which we have used in this work. The list of features present in each dataset is provided in **Table 1**. **Figure 1** provides a schematic illustration of the overall process of feature extraction and dataset construction.

**Figure 1:**
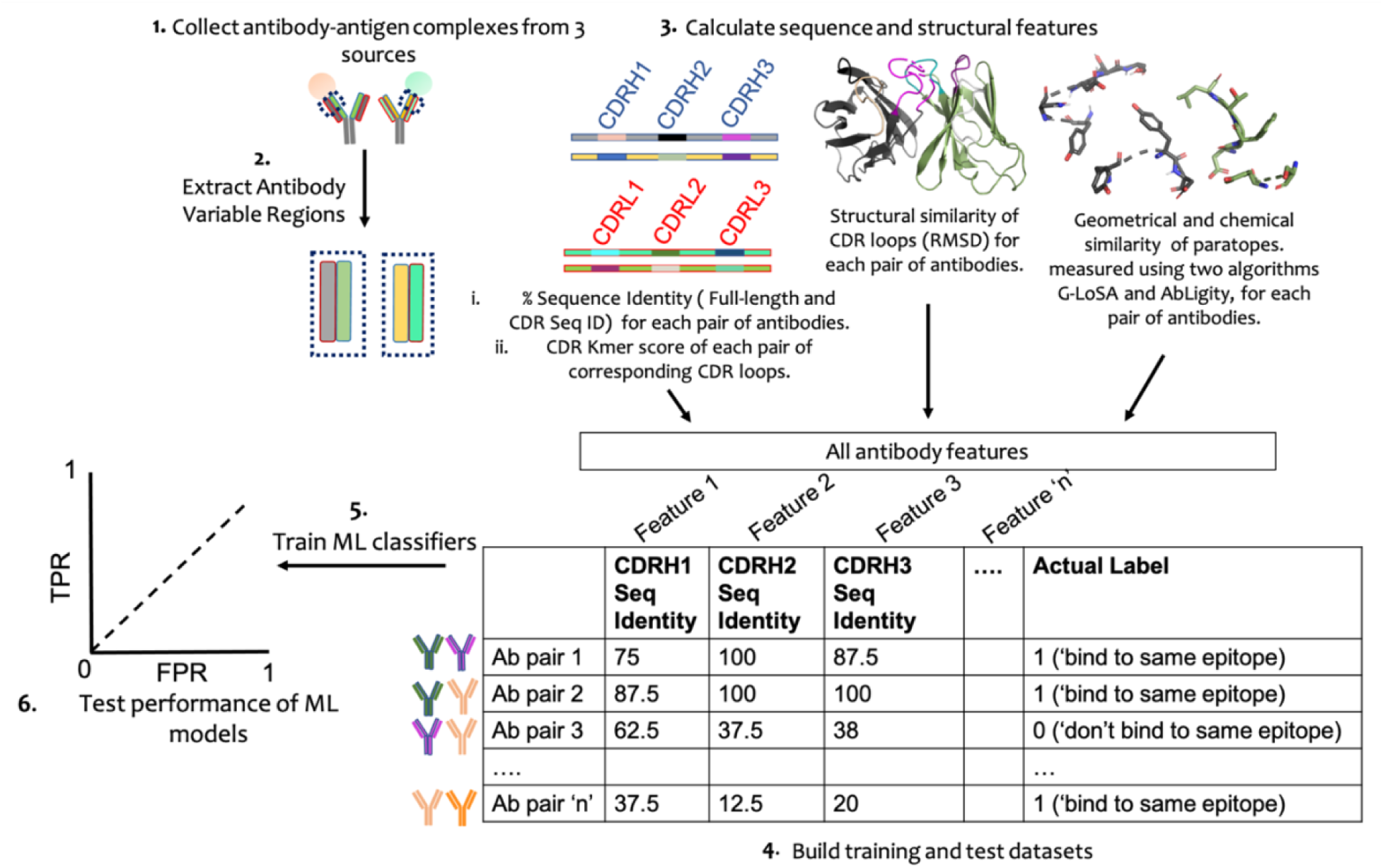
Outline of the process used to construct the datasets for training and testing the machine-learning models.

**Table 1:**
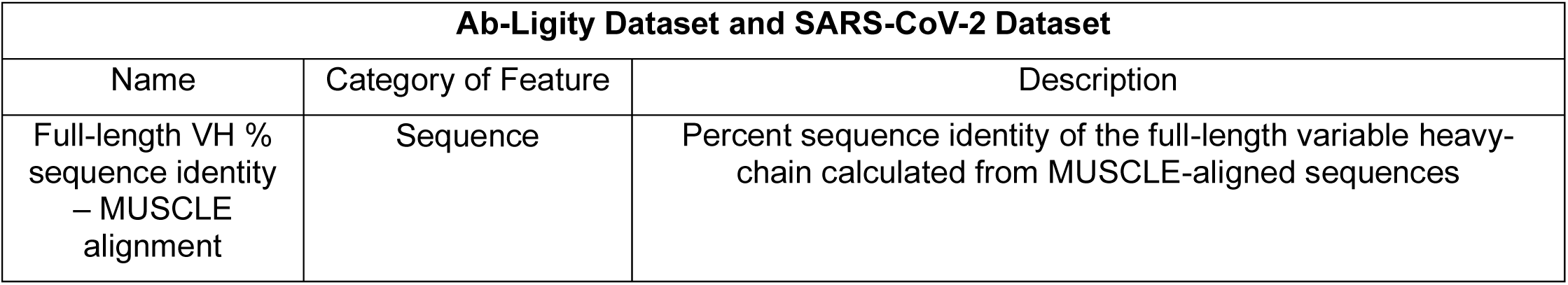
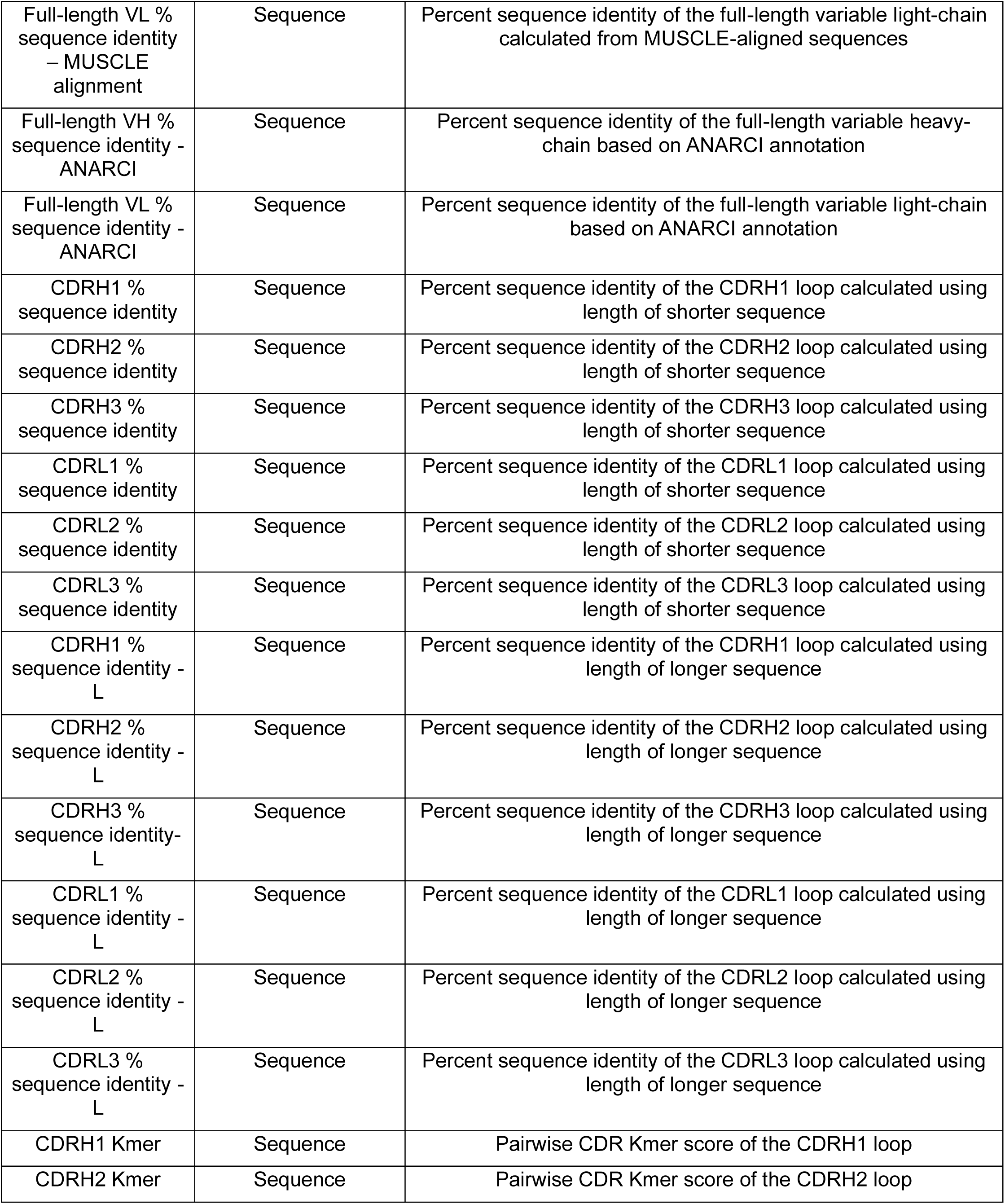
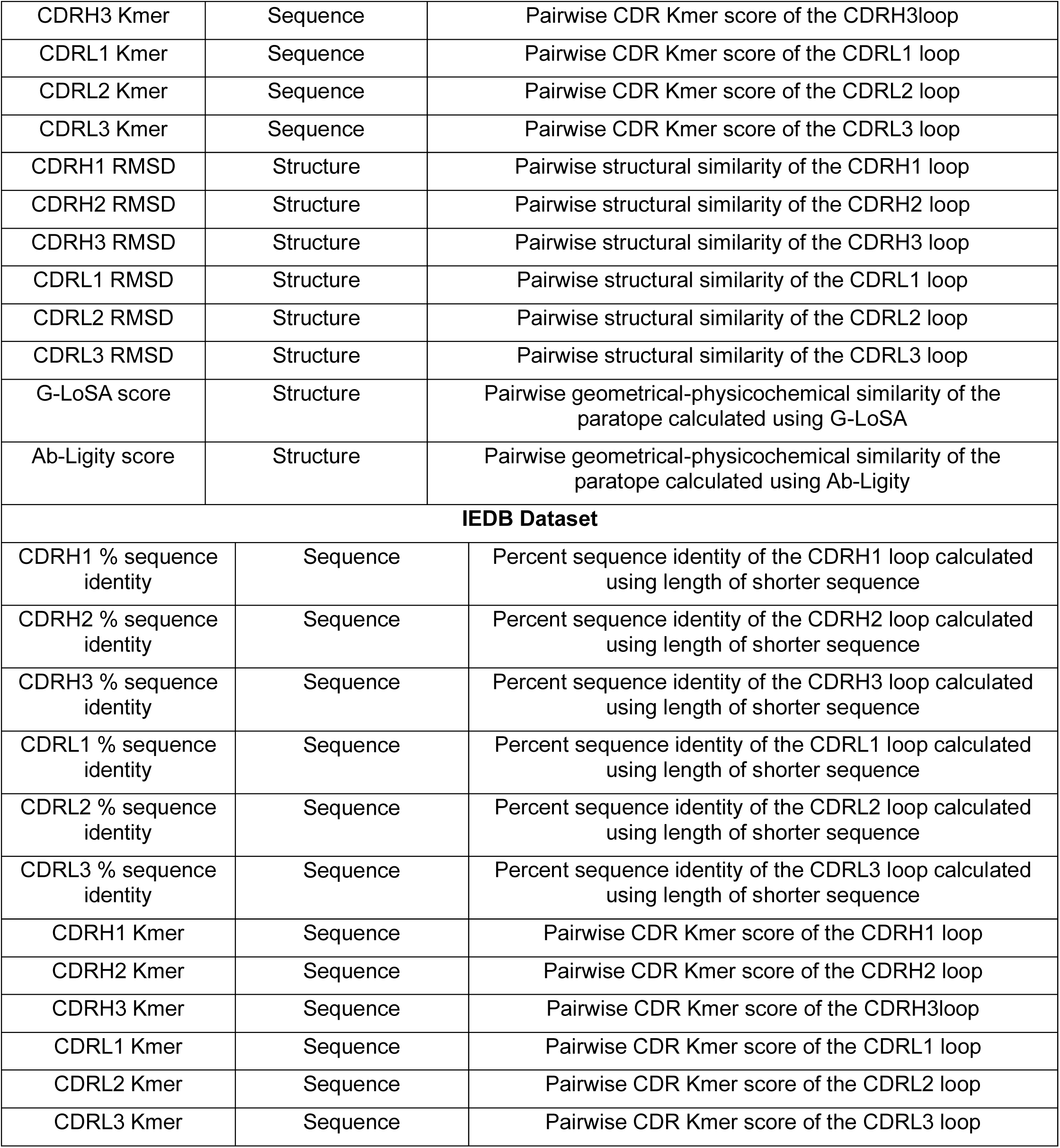
Description of antibody features present in the Ab-Ligity, SARS-CoV-2 and IEDB datasets.

We performed all-against-all comparisons of antibodies separately from each source, to estimate their similarity in sequence and structure. To compare sequence features, two types of sequence alignments were generated for the full-length VH and VL sequences using MUSCLE (*16*) and ANARCI (*17*). The MUSCLE algorithm aligns sequences of general proteins and is not restricted to antibody sequences. ANARCI is an antibody-aware algorithm as it is based on Hidden Markov Models (HMMs) of variable domain germline sequences. CDR sequences were extracted and aligned based on the ANARCI-annotated full-length VH and VL sequences. These alignments were used to calculate the pairwise sequence identity of either the full-length sequences or just the CDR sequences for every pair of antibodies derived from each source. In addition to using the identity of the CDR sequences as a metric, we also used an approach that scores a given pair of CDR sequences using a k-mer matching algorithm (*18*). Thus, there are 4 full-length sequence identity features (two pairwise sequence identity values for each of the full-length VH and VL sequences), 6 CDR sequence identity features (one sequence identity value per CDR loop), and 6 CDR k-mer scores (*18*) for each antibody pair.

Antibody pairs having 100% sequence identity over their VH and VL sequences aligned using ANARCI were removed from each dataset. Additionally, antibodies in the SARS-CoV-2 and IEDB datasets identical to antibodies in the Ab-Ligity dataset were excluded from the respective datasets to avoid redundancy. The final Ab-Ligity dataset contained a total of 334,968 unique antibody-antibody pairs out of which 333,956 pairs bind to different epitopes and 1,012 pairs bind to a common epitope. Similarly, there are 1,154 antibody-antibody pairs in the SARS-CoV-2 dataset. 910 pairs bind different epitopes and 244 pairs bind shared epitopes. Finally, the IEDB dataset consists of 174,324 antibody-antibody pairs out of which 174,039 pairs bind different epitopes and 285 bind shared epitopes.

To assess the 3D structural similarity between pairs of antibodies, we evaluated the paratope shape and the CDR loops. The geometrical similarity of the paratopes was calculated using two different binding site similarity algorithms, resulting in two similarity scores for each paratope pair. The first, G-LoSA (Graph-based Local Structural Alignment) (*19*) is a protein binding site similarity tool that employs an iterative maximum clique search algorithm and fragment superposition to identify the best alignment between two local structures. The protein structures being aligned are represented as chemical feature points, and the similarity between the two structures is quantified as a score, referred to as the GA-score, whose values range between 0 and 1. The GA-score of a random pair of structures is 0.49, while the GA-score between identical structures is 1. Ab-Ligity (*12*), is adapted from the Ligity method that uses pharmacophores for virtual screening of small molecules (*20*). It generates tokens for residues based on their physicochemical properties and employs a hashing algorithm for distance-based grouping of these residues. The outcome of this step is further used to calculate the similarity. The root mean square deviation (RMSD) metric was used to quantify the structural similarity of corresponding CDR loops between a pair of antibodies. There are 6 CDR RMSD values, one for each CDR loop, for each antibody pair.

These datasets of antibody pairs containing the respective sequence and structural features were then used to investigate the role of various features in predicting if antibodies share an epitope.

### Evaluating the predictive ability of individual features

To assess how well different sequence and/or structural features can predict whether a pair of antibodies binds to the same epitope, we split the Ab-Ligity full dataset into a training (“Ab-Ligity Full”) and test set in the ratio 3:1 (see methods) while the entire SARS-CoV-2 dataset was used as a second test dataset. For each feature, we calculated the area under the receiver operator characteristic curve (ROC-AUC) of each dataset on 1,000 simulated copies of the test dataset generated by bootstrapping with replacement to estimate a 90% confidence interval of the true ROC-AUC. To emphasize the performance of the prediction at low false positive rates, we also calculated the partial AUC capped at maximum false positive rate of 10% (AUC0.1) (*21*).

We find that the top-ranking features for the Ab-Ligity dataset include the G-LoSA score (mean ROC-AUC=0.920, AUC0.1=0.859), Ab-Ligity score (ROC-AUC=0.884, AUC0.1=0.821), features associated with the light-chain CDR3 loop (CDRL3) (CDRL3 k-mer score, CDRL3 sequence identity, CDRL3 RMSD) (**Figs. 2a,b** red arrows). The heavy-chain CDR2 loop (CDRH2) sequence features also perform closely to the CDRL3 sequence features (**Figs. 2a,b**). For the SARS-CoV-2 dataset, the CDRH2 k-mer score and sequence identity were the best predictors, with mean ROC-AUC values of 0.916 and 0.901, respectively (AUC0.1 = 0.826 and 0.819, **Figs. 2c,d**). This was followed by the full-length heavy-chain variable domain (VH) sequence identity (AUC=0.883), and the G-LoSA (AUC=0.868) and Ab-Ligity scores (AUC=0.879) (**Figs. 2c,d**). It is worth noting that for both datasets, the geometrical similarity of the paratope shape as defined by the G-LoSA or Ab-Ligity scores consistently shows a higher predictive performance, compared to that of the individual CDR structural features. Thus, independent of the CDR sequence, the shape of the paratope can also act as a good predictor of antibodies targeting similar epitopes.

**Figure 2:**
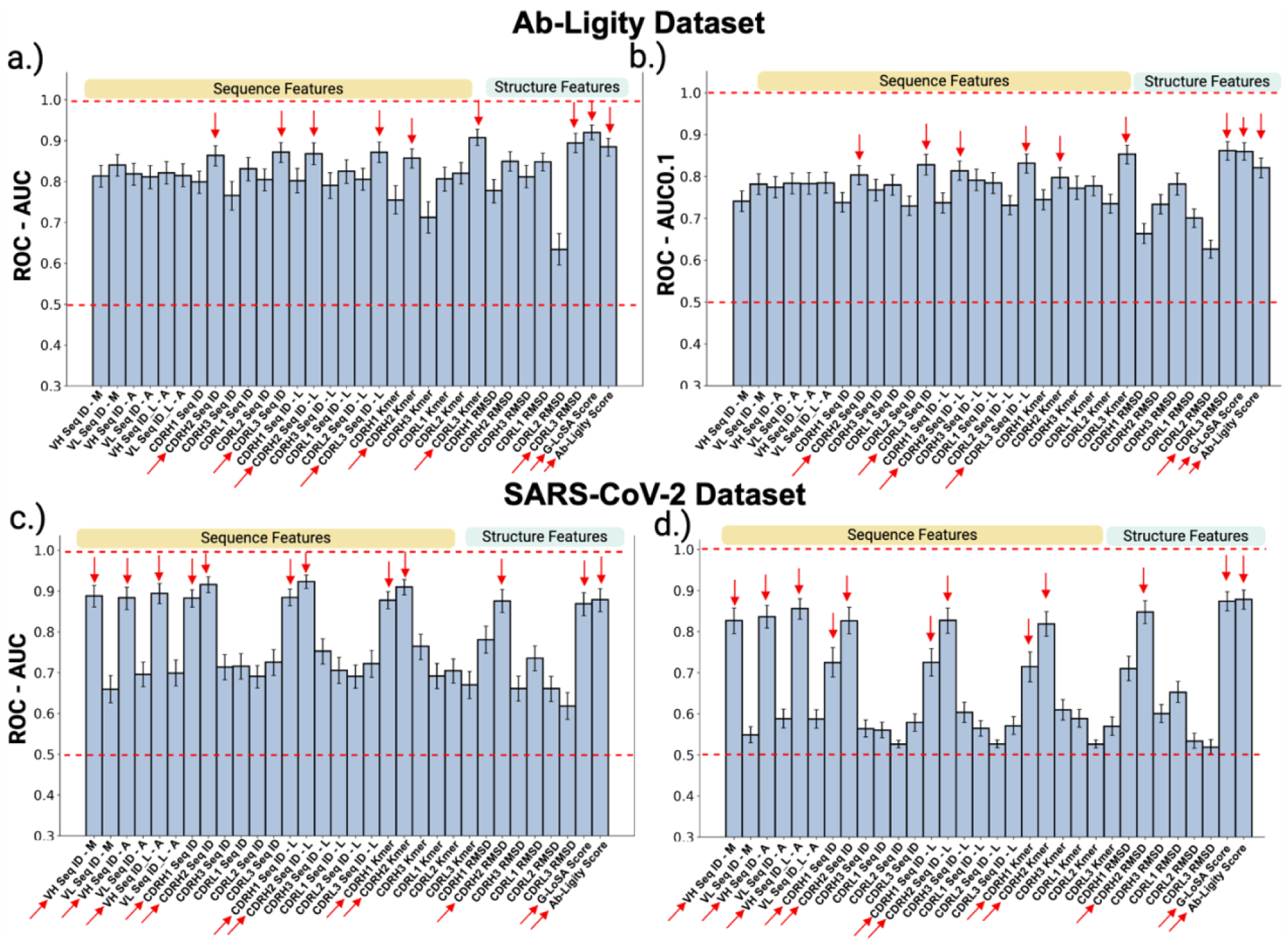
ROC-AUC and ROC-AUC0.1 values when using individual features to predict antibodies targeting a common epitope in the Ab-Ligity test dataset **(a, b)**, and the SARS-CoV-2 dataset **(c, d)**. Error bars represent the 90% confidence interval (CI). Red arrows indicate top-ranking features.L: the length of the longer sequence was used for calculating the sequence identity.

### Machine-learning models identify subsets of features important for predicting antibodies sharing an epitope

Next, we wanted to gauge whether combining two or more features would improve the prediction of antibodies binding to the same epitope. To achieve this, we trained five different machine-learning classifiers: Random Forest (RF), Logistic Regression (LR), Gaussian Naive Bayes (GNB), XGBoost (XGB) and a Feed Forward Neural Network (FFNN) with four hidden layers on the training subset of the Ab-Ligity full dataset (“Ab-Ligity Full”) (**Fig. 3a**), using a strategy in which we gradually increased the types and numbers of features included in the prediction in order to determine how important they were for successful training. Normalization of features was done by setting their mean and variance values to 0 and 1, respectively. 1,000 bootstrap samples with replacement were generated for the Ab-Ligity and SARS-CoV-2 test datasets (**Fig. 3a**) to calculate 90% confidence interval (CI) of the true AUC for each model.

**Figure 3:**
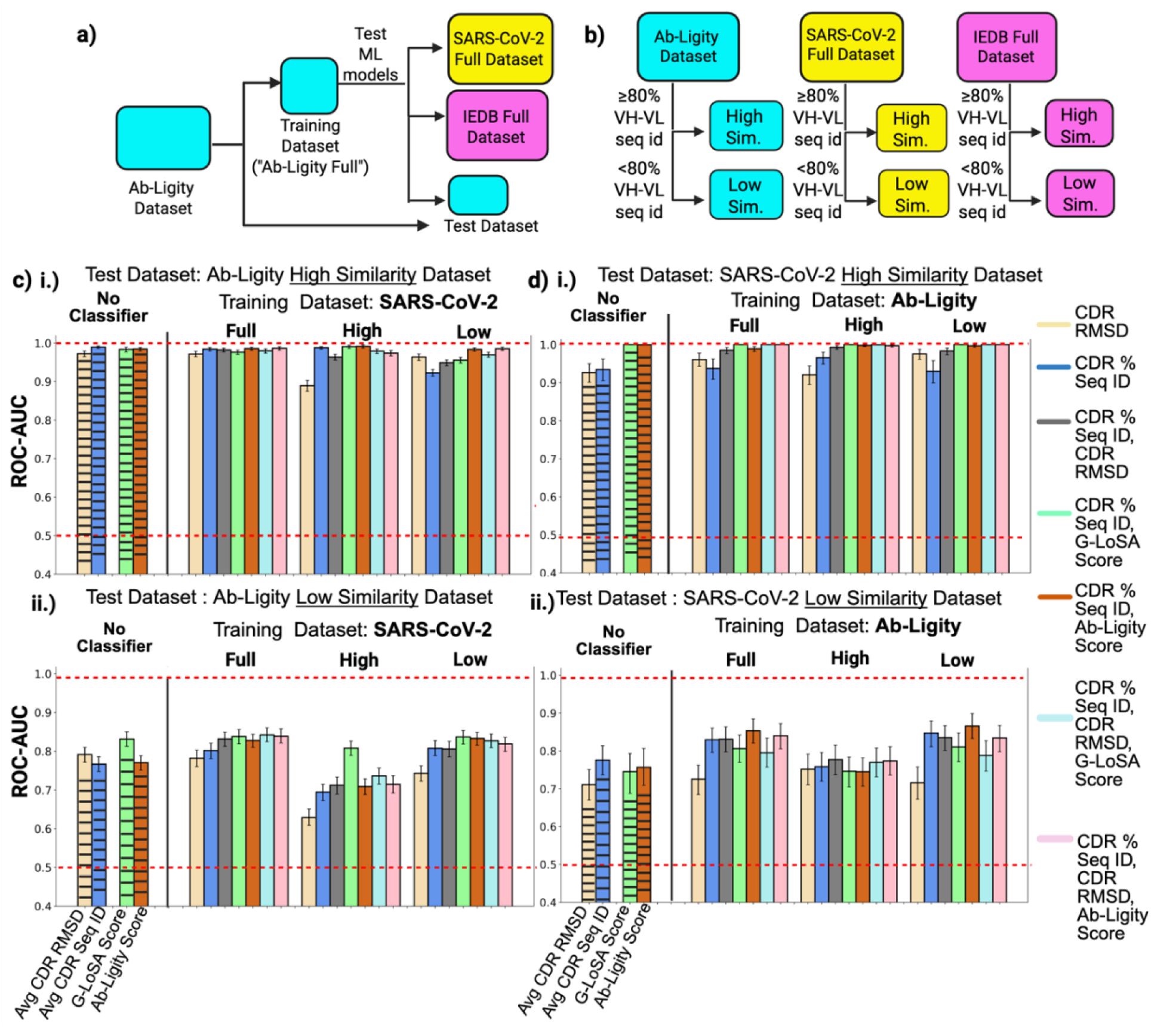
**a)**. The Ab-Ligity dataset was split into the training and test dataset. The training dataset was used to train five machine-learning models, whose performance was evaluated on the test subset, as well as two external datasets – the SARS-CoV-2 full dataset, and IEDB full dataset. **b.**) Each dataset was then split into “high-similarity” and “low-similarity” datasets using a sequence identity cutoff of 80% averaged over the VH and VL sequences. **c.)** At high levels of sequence identity, Logistic Regression models trained only on sequence features show comparable performance to models trained using a combination of sequence and structural features ((i).top panel,). Structural features, mainly represented by the binding site similarity (G-LoSA or Ab-Ligity score), lead to performance gain over models trained only on sequence features, when tested on the Ab-Ligity low-similarity dataset ((ii).bottom panel). Error bars represent the 90% confidence interval. The “No Classifier” column represents the performance of the respective features when used directly as predictors on the test dataset. Avg CDR RMSD is the average RMSD of the six CDR loops while Avg CDR Seq ID is the average percent sequence identity of the six CDR loops. **d.)** Models that learn from both structural and sequence features outperform models trained on sequence features alone, when evaluated on the SARS-CoV-2 high-similarity dataset ((i).top panel). Inclusion of structural features does not improve performance of models trained on sequence-only features when tested on the SARS-CoV-2 low-similarity dataset ((ii).bottom panel). Error bars denote the 90% confidence interval.

When assessing performance on the Ab-Ligity test dataset, we found that performance of the RF model using only pairwise sequence identity of the six CDR loops (6 features) was similar to that obtained using only pairwise RMSD of the CDR loops (6 features) (**Fig. S1a,S1b**, AUC=0.943, AUC0.1=0.929 for only CDR sequence features; AUC = 0.951, AUC0.1=0.940 for only CDR RMSD features). This pattern was observed across the other four ML models as well (**Fig. S1a,S1b**). Training the models on a combined set of CDR sequence identity and CDR RMSD features (12 features in total) resulted in a RF AUC of 0.952 and AUC0.1 of 0.944 (**Fig. S1a,S1b**). Substituting the CDR RMSD features by either the G-LoSA score or Ab-Ligity score, which quantifies the similarity between the shapes of the paratopes, negligibly improved the performances of the models (RF AUC=0.952, AUC0.1=0.929 for CDR sequence identity and G-LoSA score; RF AUC=0.956, AUC0.1=0.932 for CDR sequence identity and Ab-Ligity score, **Fig. S1a,S1b**). Next, we trained the models on the entire set of 13 features (CDR sequence identity, CDR RMSD, and either the G-LoSA score or Ab-Ligity score). The top performing models (RF and XGB) showed a marginal improvement (AUC=0.966, AUC0.1=0.951) over the corresponding models trained on combined features of CDR sequence identity and paratope shape similarity (G-LoSA/Ab-Ligity scores) (AUC=0.956) (**Fig. S1a,S1b**). Thus, these results demonstrate that the CDR sequences retain most of the information about antibody specificity for the present task.

Testing on the SARS-CoV-2 dataset indicated that models trained on only CDR sequence identity features (RF AUC = 0.898, AUC0.1=0.763) outperformed models trained on CDR RMSD features (RF AUC = 0.811, AUC0.1=0.761, **Fig. S1c,S1d**). However, it should be noted that, unlike the Ab-Ligity dataset, the SARS-CoV-2 dataset includes antibody pairs where the CDR3 loops are not of the same length. Therefore, the lower performance of CDR RMSD features in comparison to CDR sequence identity features might result from the RMSD being a poor measure for structural similarity between sequences of different lengths. When substituting the CDR RMSD by other structural features such as the G-LoSA or Ab-Ligity score, which represents the paratope shape similarity, performance was improved. A combination of CDR sequence identity and paratope shape similarity features resulted in RF AUC and AUC0.1 values of 0.920 and 0.862, respectively (**Fig. S1c,S1d**). It is possible that since G-LoSA and Ab-Ligity encode the overall geometrical structure of the paratope, they capture information about antibody specificity that might be excluded from the structure of the CDR loops. Together these results further underscore the importance of CDR sequence identity and paratope shape for predicting if antibodies share a common epitope.

### Structural features improve prediction performance when antibodies have low sequence similarity

Given that antibodies with highly similar sequences are likely to adopt highly similar structures, it is possible that the benefit of including structural information in predictions is more pronounced for antibodies that have lower sequence similarity (*12*), (*33*, *34*). We examined this using two approaches. First, we investigated the impact of the sequence similarity between the Ab-Ligity training and test data which is described in more detail in the supplementary text. Second, we split the Ab-Ligity and SARS-CoV-2 datasets each into two subsets of “high-similarity” and “low-similarity” antibody pairs (**Fig. 3b**) using a cutoff of 80% sequence identity averaged over the full-length heavy (VH) and light-chain (VL) variable domain sequences. The cutoff was chosen so that the number of antibody pairs in the positive class was approximately similar in both the high-similarity and low-similarity subsets of the Ab-Ligity and SARS-CoV-2 datasets. The details of the resulting subsets are listed in **Table 2**.

**Table 2:** Details of antibody pairs present in the high-similarity and low-similarity subsets of each dataset.

|  | **Total number of antibody pairs in dataset** | **No. of antibody pairs in the positive class** | **No. of antibody pairs in the negative class** |
| --- | --- | --- | --- |
|  | <u>Ab-Ligity Dataset</u> |  |  |
| High-similarity | 2942 | 516 | 2426 |
| Low-similarity | 332026 | 496 | 331530 |
|  | <u>SARS-CoV-2 Dataset</u> |  |  |
| High-similarity | 260 | 130 | 130 |
| Low-similarity | 894 | 114 | 780 |
|  | <u>IEDB Dataset</u> |  |  |
| High-similarity | 618 | 73 | 545 |
| Low-similarity | 173706 | 212 | 173494 |

We used 12 different train-test combinations along with bootstrapping to evaluate model performance (**Fig. S2**). Models were trained on one of the following: the full SARS-CoV-2 dataset, the SARS-CoV-2 high-similarity dataset, and the SARS-CoV-2 low-similarity dataset (**Fig. S2a**). Evaluation of these trained models on the Ab-Ligity high-similarity dataset showed that irrespective of the type of training dataset used (full, high-similarity, or low-similarity), sequence identity features alone could strongly predict if two antibodies share an epitope and including structural features did not have any advantage (**Fig 3c.i**; AUC of striped blue bar=0.989, **Fig. S3a**). In contrast, when the models were tested on the Ab-Ligity low-similarity dataset, structural features became important for improving predictions. Moreover, models trained on only the low-similarity subset of the SARS-CoV-2 dataset showed comparable performances to the corresponding models trained on the full dataset. For these models, combinations of CDR sequence identity and the G-LoSA score had the highest predictive performance (LR AUC=0.838, AUC0.1=0.753,) followed by CDR sequence identity combined with either CDR RMSD (LR AUC=0.831, AUC0.1=0.754.) or Ab-Ligity score (AUC=0.827, AUC0.1=0.706) (**Fig 3c.ii**, **Fig. S3c,S3f)**. The models that utilized only CDR sequence identity features underperformed. The AUC and AUC0.1 values of the other ML models are shown in **Figs. S4-S7**. Models trained on only the high-similarity dataset demonstrated substantial performance loss across all feature combinations on the low-similarity test dataset, when compared with models trained on the full or low-similarity datasets (**Fig. 3c.ii**). Within this category, models that utilized a combination of CDR sequence identity and G-LoSA score had the highest performance (**Fig. 3c.ii**, LR AUC=0.807,AUC0.1=0.695, **Figs S3d,S3f**). Overall, these results highlight that structural features become more informative when the sequence identity between two antibodies is low. Including antibody pairs with low sequence identity in the training dataset is thus vital for reliable model performance.

We then repeated this process by swapping the training and test datasets. Models were trained on the full, high- and low-similarity subsets of the Ab-Ligity dataset and tested on the SARS-CoV-2 datasets (**Fig. S2b**). As expected, the sequence similarity of antibody pairs in the training dataset again did not make a difference when the models were evaluated on the high-similarity SARS-CoV-2 dataset. When models were tested on the low-similarity SARS-CoV-2 dataset, models which were trained on the full- or low-similarity datasets performed better than those trained on high-similarity datasets alone. Models utilizing only CDR sequence identity features had similar performance to models that used both sequence identity and structural features (**Fig. 3d.ii.**, **Figs. S8-S11**). For example, the LR models that used only CDR sequence identity features in the full or low-similarity training datasets had AUC values of 0.829 and 0.847, respectively (**Fig.3d.ii.**). Addition of the G-LoSA or Ab-Ligity scores did not improve their performances. (**Fig.3d.ii.**). This contrasts with what we observed with models evaluated on the Ab-Ligity low-similarity dataset. This suggests that despite the models learning from structural features in the training dataset, the structural features in the SARS-CoV-2 test dataset do not provide additional information beyond those encoded in the sequence. This could be due to the SARS-CoV-2 dataset being comprised of antibodies obtained early in the Covid19 pandemic, which is expected to consist of antibodies that were generated after a single priming, resulting in low levels of somatic hypermutation (SHM) and their sequences being very close to the germline sequences, as demonstrated in other studies (*24*).This could result in the antibodies having very similar structures with low structural variation which limits the ability of structural features to distinguish between antibodies binding different epitopes.

### Testing on IEDB dataset reveals importance of having the right training dataset

The Immune Epitope Database (IEDB) (*25*) contains experimentally validated B cell epitopes and sequences of the antigen receptors recognizing them, if available. We saw earlier that the five ML models showed good performances on the Ab-Ligity test dataset and the SARS-CoV-2 dataset, even when only CDR sequence identity features were used. Therefore, we wanted to test how well these models perform on the IEDB dataset if we restricted the features to only CDR sequence identities. We first trained the five ML models on the training component of the full Ab-Ligity dataset (“Ab-Ligity Full”) taking into account only the sequence identities of the six CDR loops, and tested their predictive performance on the full IEDB dataset. All five models showed similar mean AUC values between 0.786 and 0.802 (**Fig. S12**).

Next, we split the IEDB dataset into the “high-similarity” and “low-similarity” subsets using an average sequence identity of 80% over the VH and VL sequences. The number of pairs in the positive and negative classes is listed in Table 2. The five ML models were each trained separately on three datasets: Ab-Ligity Full, Ab-Ligity high-similarity dataset, and Ab-Ligity low-similarity dataset. The performance of each trained model was tested on the IEDB high-similarity dataset (**Fig. 4a, Fig. S13a**), and the IEDB low-similarity dataset (**Fig. 4b, Fig. S13b**). As expected, all models showed higher predictive performance (AUC ranges between 0.884 to 0.911) on the high-similarity test dataset than on the low-similarity test dataset (range of mean AUC values:0.546 to 0.717). For the low-similarity test dataset, we again noticed that the models trained on the high-similarity dataset performed worse than models trained on the full or low-similarity datasets (**Fig. 4b, Fig. S13b**). This observation further reinforces the importance of having antibody pairs with diverse sequence identities in the training dataset for model performance.

**Figure 4:**
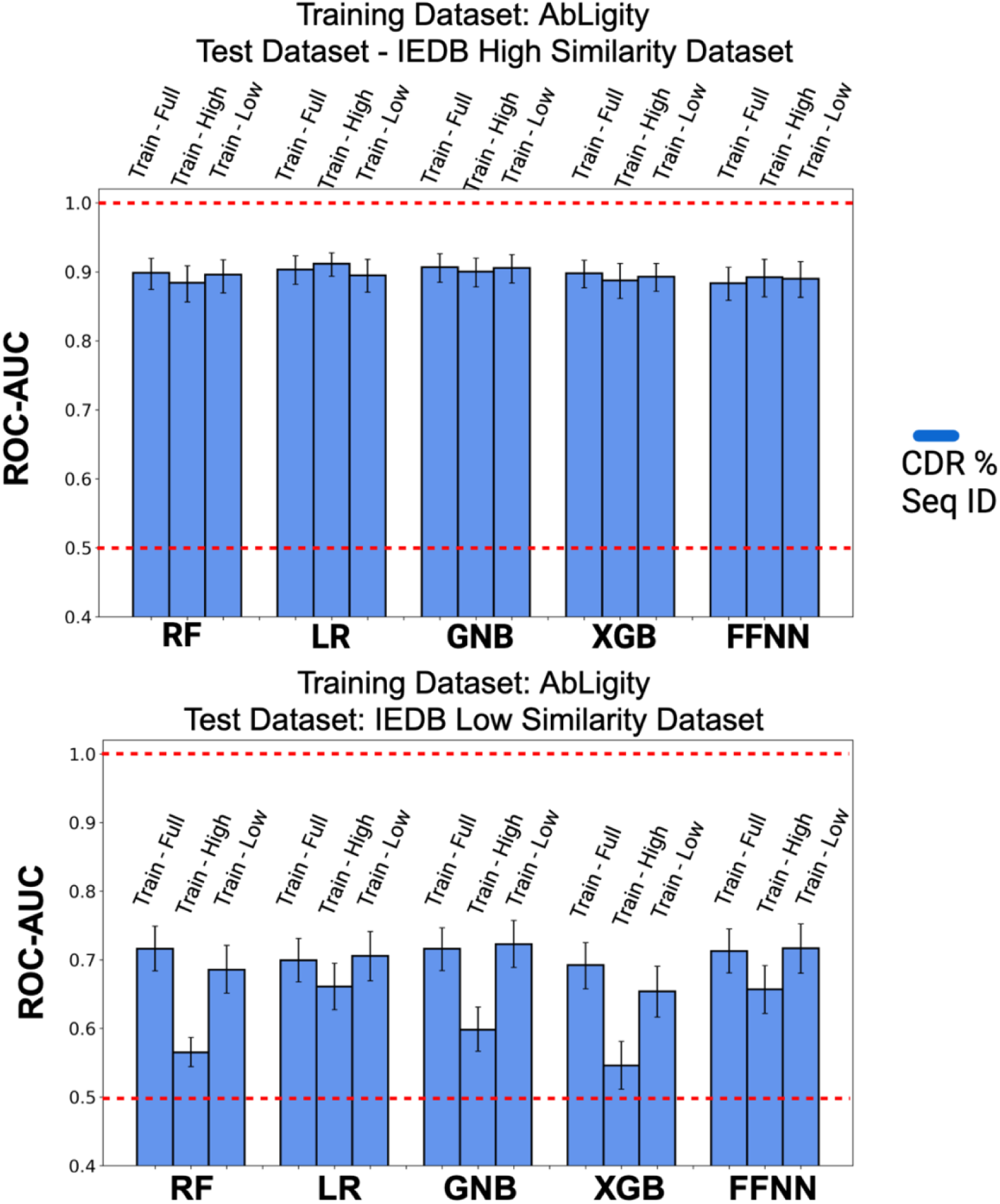
Training on low sequence identity pairs leads to better model performance on the IEDB low-similarity dataset. ROC-AUC values of the different ML models when tested on the IEDB high-similarity dataset (top) and the IEDB low-similarity dataset (bottom). Error bars represent the 90% confidence interval.

### Replacing sequence identity score metric with k-mer similarity score metric does not substantially impact model performance

Our group previously developed a method to evaluate the similarity of TCR sequences (*18*) using a “k-mer similarity score”, to quantify the similarity between pairs of TCR sequences without the need of aligning sequences. We investigated whether substituting aligned CDR sequence identity with their corresponding k-mer similarity scores would affect the performance of the ML models. We trained models on either the full Ab-Ligity training dataset or the full SARS-CoV-2 dataset, using CDR k-mer scores instead of CDR sequence identities (**Fig 5, Fig. S14**). Evaluation on the full test datasets (Ab-Ligity, SARS-CoV-2, and IEDB) showed that using the CDR k-mer score in place of the CDR sequence identity feature did not substantially influence the performance of models trained on equivalent feature combinations (**Fig. S14**). When the low-similarity datasets were used for testing, we observed a slight reduction in model performance after substituting the CDR sequence identities by the CDR k-mer scores (**Fig 5**). For instance, the LR model trained on the SARS-CoV-2 dataset and tested on the Ab-Ligity low-similarity dataset had an AUC of 0.706 when only CDR k-mer scores were used (**Fig. 5a**), and an AUC of 0.802 if only CDR sequence identities were used (**Fig 3c.ii.**, “Full”). Testing on the IEDB low-similarity dataset revealed no major differences in performance for the LR model when interchanging between CDR k-mer score features (**Fig. 5c**) and CDR sequence identity features (**Fig. 4, lower panel**). The other models also displayed a similar trend (**Figs. S15-17**). Overall, these results show that the CDR k-mer score can be used instead of CDR sequence identity, as no sequence alignment is required, which is beneficial if no full-length sequences are available.

**Figure 5:**
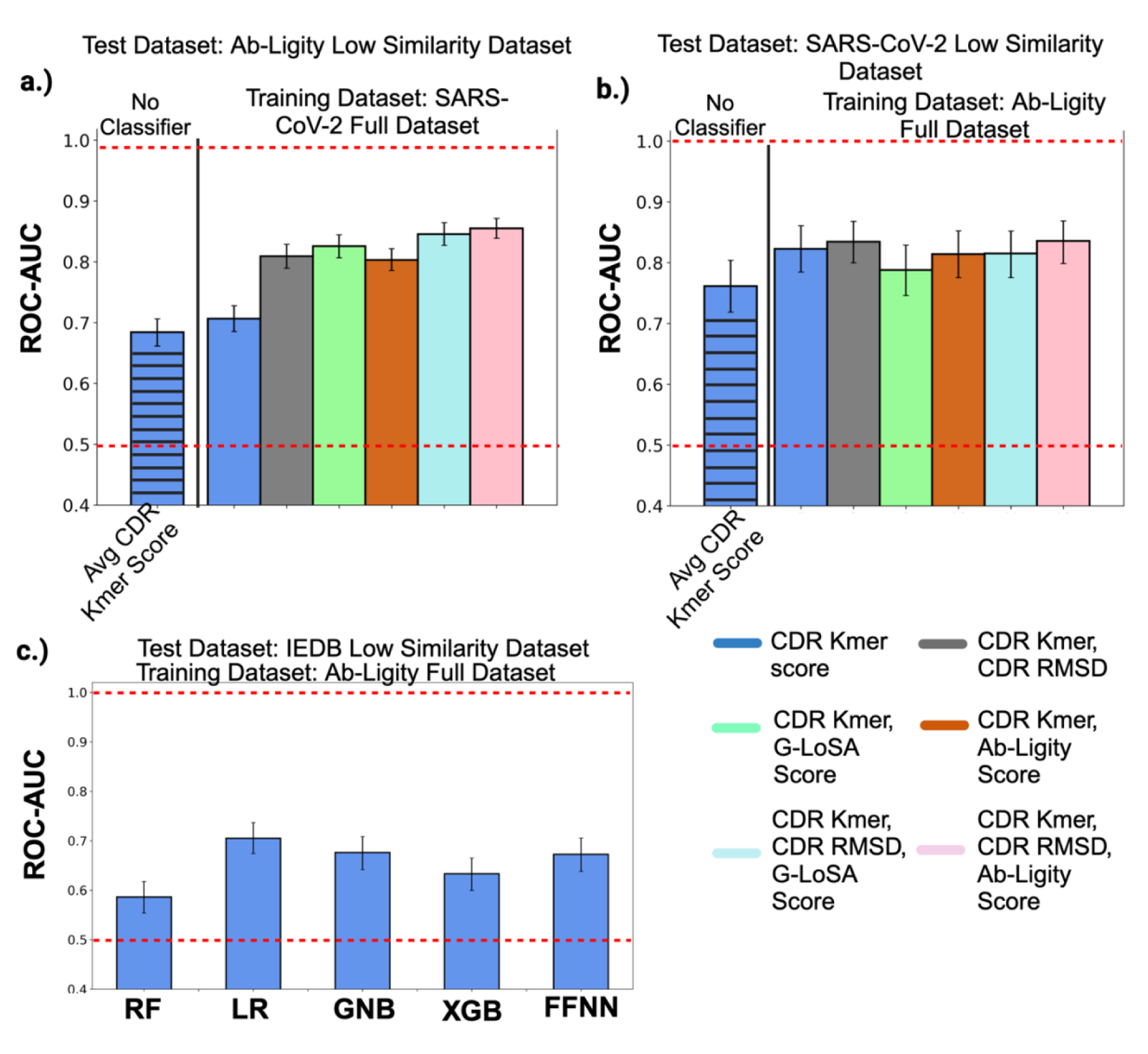
Replacing the CDR sequence identity by its corresponding k-mer score does not lead to major differences in model performance. **a.)** ROC-AUC values of the LR models tested on the Ab-Ligity low-similarity dataset (trained on the SARS-CoV-2 full dataset). The “No Classifier” column indicates the performance of the average CDR k-mer score when it is used directly as a predictor on the test dataset. **b.)** Performance on the SARS-CoV-2 low-similarity dataset (trained on Ab-Ligity full dataset). **c.)** AUC values of the five ML models when tested on the IEDB low-similarity dataset (trained on Ab-Ligity full dataset). RF:Random Forest, LR:Logistic Regression, GNB:Gaussian Naive Bayes, XGB:XGBoost, FFNN:Feedforward Neural Network.

### The BCRMatch standalone tool

The results of this study were used to inform the design of a standalone tool termed ‘BCRMatch’ that can be freely downloaded and used as a Docker container or installed locally (https://github.com/IEDB/BCRMatch). Since we evaluated five different ML methods in this work, we also tested the performance of an ensemble obtained by combining different classifiers. Each ML classifier was used to predict the probabilities of antibody pairs in the entire IEDB and SARS-CoV-2 datasets sharing an epitope. The scores from both datasets were combined into a single dataset. Random antibody pairs were chosen from both positive and negative classes in the IEDB dataset and assigned to a test dataset. Percentile ranks were calculated for each antibody pair in this test dataset, using the distribution of scores previously calculated on the IEDB and SARS-CoV-2 datasets. Following this, we calculated the harmonic mean of the percentile ranks of the five classifiers, as well as only that of the top-ranking classifiers (LR and GNB), for each antibody pair. The ROC-AUC values of the individual classifiers’ percentile ranks and the harmonic mean of the percentile ranks are shown in **Fig. S18**. Based on these findings, we define the BCRMatch tool as an ensemble of the LR and GNB classifiers. The tool can accept either the full-length VH and VL sequences, or the sequences of the six CDR loops of antibodies as input. The tool extracts the CDR sequences if VH and VL sequences are provided. It then calculates the k-mer scores of corresponding CDR sequences for each antibody pair, and uses the pre-trained machine-learning models to make the predictions. The output of the tool is a table containing the probability values of the LR and GNB classifiers, their corresponding percentile ranks, and the harmonic mean of the percentile rank, for each antibody pair. In the future, we plan to build a BCRMatch webserver that will be hosted on the IEDB tools website.

### Examining the impact of gene usage vs. CDRs

We were intrigued by the high performance of the CDRH2 sequence identity feature on both the Ab-Ligity and SARS-CoV-2 datasets when it is used directly as a predictor, and in particular how it outperformed the CDRH3 performance. To confirm this finding, we repeated this step on the IEDB dataset and observed that the CDRH1 and CDRH2 sequence identity features also had better performance than CDRH3 sequence identity (**Fig. 6** and **Fig. S19**). Since, the CDRH1 and CDRH2 sequences are encoded by the V germline genes, we asked if predicting shared epitopes could simply be done by identifying common V genes. We used ANARCI to assign the closest-matching V germline gene to each of the heavy-chain and light-chain sequences of the antibodies. Antibody pairs that had the same heavy-chain or light-chain V gene were assigned a score of 1 while pairs that have different V genes were assigned a score of 0. A score of 2 was assigned to pairs that have both same heavy-chain and light-chain V genes. The performances of these three V gene features as predictors were evaluated in the same manner as the other individual features, on the Ab-Ligity and IEDB datasets. Remarkably, the V gene features showed high performance on both datasets (**Fig. 6**) indicating that many antibodies binding a common epitope also share the same V genes. However, the performance of CDRH1 and CDRH2 sequence identity features still outperformed the V gene features (**Fig. 6**). This strongly indicates that the CDRH1 and CDRH2 sequences are of vital importance, possibly more than CDRH3, in predicting antibody specificity.

**Figure 6.**
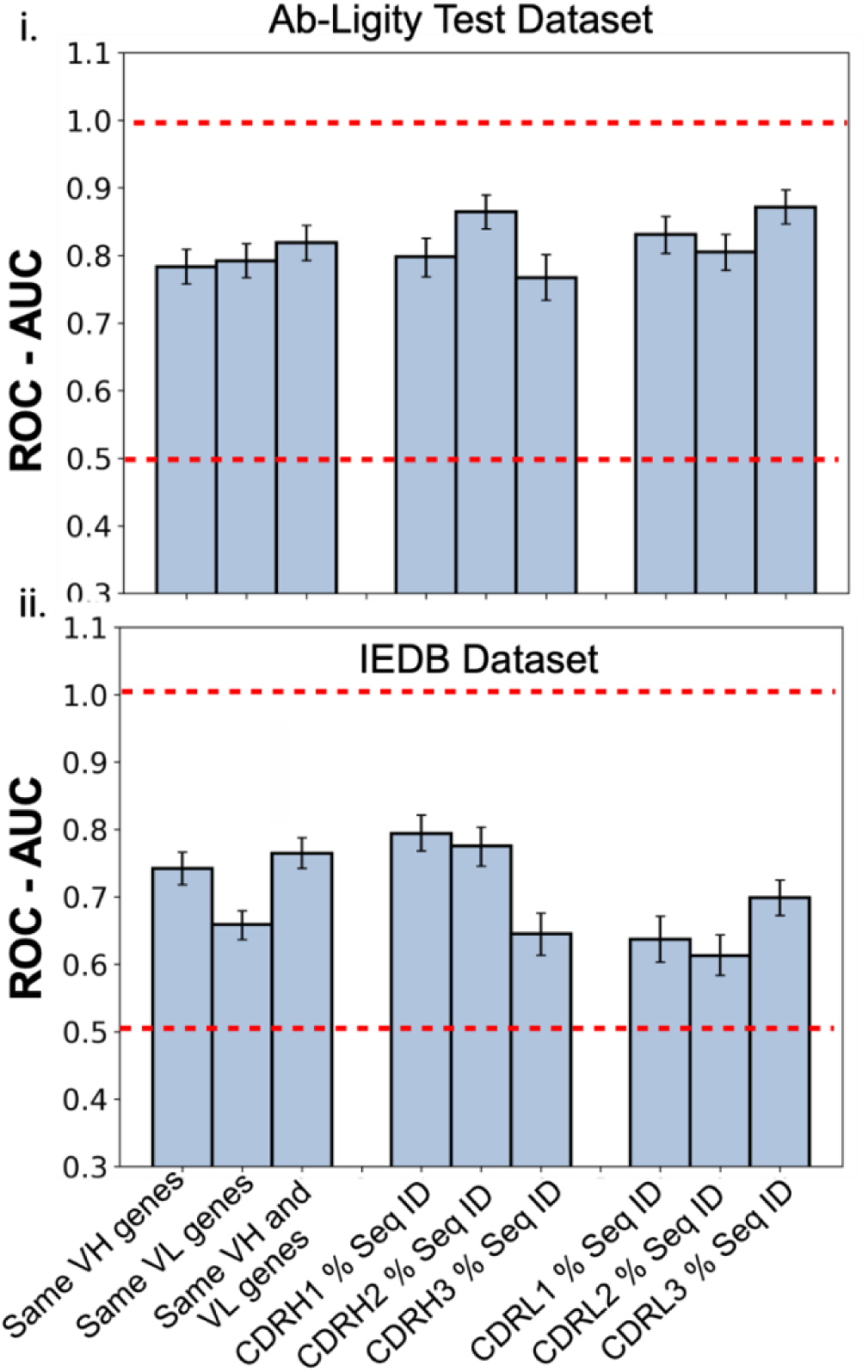
Same V genes are highly predictive of antibodies sharing an epitope but not as much as the CDR1 and CDR2 sequences. Comparison of ROC-AUC values when using same heavy-chain V gene, light-chain V-gene, or both, as predictors, alongside CDR sequence identity in the i. Ab-Ligity test dataset. ii. IEDB dataset.

## DISCUSSION

Many machine-learning tools for predicting antibody specificity, antibody epitopes, or antigen-antibody interactions exist (*36*-*39*). These tools adopt various methodologies such as training on only antibody sequences (*28*), on sequences of antibodies specific to a particular antigen such as the SARS-CoV-2 Spike or influenza HA proteins (*41*,*42*), on heavy-chain variable domain sequences (*31*), or on structures of antibody-antigen complexes (*44*-(*33*)*46*). Fewer studies have described and compared clustering methods that use both sequence and structural information to predict groups of antibodies common to an epitope. Xu and colleagues trained a support vector machine (SVM) classifier on sequence and structure-based features of pairs of BCRs and used this to derive a similarity score (*35*). This similarity score was then used to hierarchically cluster the BCRs. The authors also state that the information in the classifier comes mainly from sequence features although structural features also help improve classifier accuracy. They further build on that work by developing a method that clusters antibodies based on only CDR sequences of the heavy chain variable region (CDRH) (*36*). Another study by Chomicz and colleagues performed a benchmarking of different clustering methods for antibodies and created an online tool CLAP that enables users to visualize clusters grouped using the different clustering methods (*37*). Our study here expands on this previous work by specifically investigating the contribution of different features of the antibodies to their overlapping binding specificity while controlling for high vs. low-sequence similarity.

An interesting finding from this study was how the model performances on the low-similarity test datasets varied with the type of training data used. When the ML models were trained on the high-similarity datasets, using only CDR sequence identity features, they showed reduced performance on all low-similarity test datasets. Their performance on the low-similarity test datasets improved when either a mix of both high- and low-similarity, or only low-similarity antibody pairs were used to train them. It also became clear that when only high-similarity antibody pairs were used to train the ML models, the contribution of structural features greatly improved model performance on the low-similarity test datasets. This demonstrates that including antibody pairs with a wide range of sequence similarities in the training dataset can boost model performance.

A surprising revelation was how the CDRH1 and CDRH2 sequence features consistently outperformed CDRH3 sequence features and V gene identity. While antibody pairs that bind a common epitope share the same V genes in the majority of cases, pairs having different heavy-chain or light-chain V genes binding the same epitope were also present in our datasets. This prompted us to investigate if these V gene pairs occurred more frequently than what would be expected by chance, since they would need to share similar sequence or structural features that enable them to bind the same epitope. We calculated the observed and expected frequency of the co-occurrences of the V gene pairs, as well as the fold change of observed over expected frequency. The fold change values are plotted as a heatmap in **Fig. S20a, S20b**. The heavy-chain V gene IGHV1-46 co-occurred often with IGHV1-2 and IGHV1-18. These genes were present mainly in antibodies that target the CD4 binding site on HIV gp120 (*38*). The high similarity between the CDRH2 sequences of these antibodies made it hard to distinguish whether they originated from IGHV1-2 or IGHV1-18 germline genes. Among gene pairs in the IEDB dataset, heavy-chain genes IGHV7-1 and IGHV3-1 had the highest fold change, followed by the gene pair IGHV3-33 and IGHV1-2. Upon closer inspection of the antibody pairs which were encoded by the frequently co-occurring heavy-chain V genes (IGHV1-46 and IGHV1-18, IGHV7-1 and IGHV3-1, IGHV3-33 and IGHV1-2), we observed shared motifs in the CDRH2 and CDRH1 sequences of these genes (**Fig. S20c**).

Our findings here are supported by previous studies which showed that antibodies use germline-encoded amino acid binding (GRAB) motifs to recognize public epitopes (*39*). In their study, the authors discovered lysine-specific GRAB motifs and suggest that these motifs could also have specificity for certain amino acid combinations. Further extensive work will be needed to discover additional motifs. Findings from this study and our study both suggest a mechanism in which the CDR1 and CDR2 regions are involved in initial recognition of the epitope while the CDR3 region contributes to increasing the affinity between the antibody and antigen without changing the epitope. This highlights the importance of the CDRH1 and H2 loops in mediating antibody specificity to epitopes.

A highly cited study for the importance of CDRH3 for antibody epitope specificity was published by Xu and Davis (*40*). In their study, the authors use transgenic mice harboring only a single human V_H_5-51 gene, and different D_H_ and J_H_ genes. Immunization of these mice with different antigens and sequencing of these antigen-specific antibodies derived from the immunized mice revealed that these antibodies differed in their CDRH3 sequences while the rest of the sequences remained the same. They attributed the specificity of the antigen-specific antibodies to the differences in their CDRH3 sequences. While this is true, the study does not show the relative importance of the CDR1/CDR2 versus the CDR3 loops since the CDR1/2 sequences are genetically set in this mouse model and could obtain diversity only due to somatic hypermutation. The study by Shrock and co-authors (*39*), shows that antibodies containing the IGHV5-51 gene bind to different antigens using a similar binding motif and reveal the presence of a GRAB motif in IGHV5-51. This GRAB motif can also be found in the sequences of the antibodies described in the study by Xu and Davis. Therefore, this raises a possibility that while the CDRH3 can lead to recognition of different antigens, it may not be as important as the CDRH1 and CDRH2 for conferring specificity/initial epitope recognition to the antibody when a diversity of V genes is available.

The BCRMatch tool described here is built on the insights gained from this work, and can be used for predicting epitopes of antibodies based on shared features with other antibodies having known epitopes, thereby enabling researchers to rapidly predict epitope candidates for their antibodies of interest. The tool will be made publicly available and we anticipate that it will become increasingly valuable as we continue to gather more antibodies with known specificities in the IEDB.

One caveat of this work is that the antibody pairs that do not share an epitope, which we consider as the negative data in this study, has not been experimentally validated. Another caveat is the limited number of antibodies included in the SARS-CoV-2 dataset. The dataset was built using antibodies downloaded from the Cov-AbDab in late 2020, when the original SARS-CoV-2 virus dominated the pandemic, and structures added later were not included in the dataset. However, the datasets will be updated and models re-trained on a periodic basis, to leverage the ever-expanding amount of publicly available antibody data.

## MATERIALS AND METHODS

### Antibody sources for constructing the datasets

#### IEDB dataset

Curated BCR entries were retrieved from the Immune Epitope Database (IEDB,www.iedb.org) (Vita et al., 2024) (*15*) and are provided in **Table S1**. Each entry contains information about the receptor group to which the BCR entry belongs, the receptor id, the variable heavy (V, D, J) and light-chain (V,J) genes, the full-length variable domain sequences of the heavy- and light-chains, the CDR sequences, the epitope, antigen containing the epitope, organism to which the antigen belongs, PDB ID of any available structures, and number of assays describing the specific BCR. The receptor id is a numeric IEDB identifier that is assigned to a specific immune receptor. Receptors sharing the same CDR3 sequence are grouped under a “receptor group” (*15*). The IEDB definition of a B-cell epitope requires mapping to a molecular structure of not more than 5 kiloDaltons. These must have been experimentally validated for binding to a BCR/ antibody. The residues comprising the epitope are defined in the “Epitopes” data field in the Results page after a query is performed in the search interface of the IEDB home page (*25*). Receptors without full-length variable domain sequence data were excluded. For each unique epitope entry in the dataset, which is the set of residues in the epitope not identical to another entry in the same dataset, BCRs with non-redundant receptor ids that bind the particular epitope were compiled into a list. Receptor ids listed under a specific epitope are considered to bind to the same epitope. Therefore, all possible pairs of B-cell receptor ids falling under this epitope are assigned to the positive class. Pairs of B-cell receptor ids where each B-cell receptor binds to a different epitope were included in the negative class.

#### Ab-Ligity dataset

The original dataset of antibody pairs used in the Ab-Ligity study (*12*) contains 920 antigen-antibody complexes and their corresponding Ab-Ligity scores and is restricted to only antibody pairs in which their CDRH3 loops are of equal length. Only a single copy of each antibody is present in the original Ab-Ligity dataset. For the purpose of our study, upon request to the authors, we also obtained antibody models and predicted paratopes used to test the Ab-Ligity method. In their study, the authors used ABodyBuilder (*41*) to build antibody models, and for predicting paratope residues, another method, Parapred (*42*) was utilized. Parapred assigns scores to CDR residues that reflects their likelihood of binding. Only those entries for which modelled antibody structures were available were included in the final dataset.

#### SARS-CoV-2 dataset

SARS-CoV-2 antibody-antigen complex structures were manually retrieved from the Coronavirus Antibody Database (CovAbDb) (*13*) in October 2020. Atomic coordinates of epitope residues from every complex were extracted using an in-house script. A residue on the antigen chain was defined to be part of the epitope if any of its non-hydrogen atoms were located within a 5 Å distance from any non-hydrogen atom on the antibody chain. These epitope residues were saved into separate files in PDB format. Using a protein binding site similarity algorithm (G-LoSA) (*19*), described in detail in the next section, we quantified the epitope similarity by calculating G-LoSA scores for every pair of epitopes. These scores are illustrated as a heatmap in (**Supp. Fig. 1**). Based on this and combined with visual inspection of each antibody-antigen complex structure, the epitopes were categorized into one of the epitope classes defined in (*43*) (**Fig. S21**). Class 1 overlaps with the angiotensin-converting enzyme 2 (ACE2)-binding site on the receptor-binding domain (RBD) of SARS-CoV-2 and is accessible only when the RBD is in the “up” conformation. Class 2 also overlaps with the ACE2 binding site but can be accessed when the RBD is both in the “up” or “down” conformation. Class 3 does not overlap with the ACE2 binding site and can be accessed in both the “up” and “down” conformations, while Class 4 also does not overlap with the ACE2 binding site but is accessible only in the “up” conformation. Due to the highly overlapping nature of the epitopes on the RBD, we set a stringent G-LoSA score (which translates to high epitope similarity) of 0.75 as the threshold for assigning class labels. Antibody pairs whose corresponding epitopes had a G-LoSA score of 0.75 or above were considered to belong to the positive class (binding to a common epitope), and the rest were assigned to the negative class (binding to a different epitope).

### Estimating three-dimensional (3D) similarity of paratopes

The geometrical similarity/similarity of the 3-D binding site was assessed using two methods.

#### G-LoSA

The Ab-Ligity study (*12*) contains information about predicted paratope residues on the modelled antibodies in their dataset. To calculate the paratope similarity of these paratopes using G-LoSA (*19*), the atomic coordinates of the predicted paratope residues were extracted from the pre-generated full-length 3D models of the antibodies (*12*) using in-house scripts. All-against-all pairwise comparison of the paratopes was carried out to obtain the G-LoSA scores. For the SARS-CoV-2 dataset, experimentally-determined 3D structures of the antigen-antibody complexes were used in which the CDR loops can be in a different conformation from the unbound state. A two-step process was therefore used to extract paratope residues. In the first step, residue details such as residue number, residue name and atomic coordinates of paratope residues located within 5 Å of any antigen non-hydrogen atom were extracted and saved into separate PDB files. In the second step, antibody-antigen complex structures were separated, and the side-chain residues of all antibody structures were subjected to conformational rearrangement using the SCWRL4 algorithm to mimic the unbound state of the antibody. Using the details of residue ids obtained in the first step, the modified atomic coordinates of the paratope residues were extracted from the SCRWL4 processed antibody structures and saved into separate files in PDB format. These files were used for all-against-all G-LoSA similarity comparison of the paratopes in the SARS-CoV-2 dataset.

#### Ab-Ligity

In addition to G-LoSA, we also used Ab-Ligity (*12*) to measure the paratope similarity. The Ab-Ligity algorithm tokenizes both paratope and epitope residues according to their chemical properties (aliphatic, aromatic, basic, acidic to name a few) and then uses a hashing function to bin these tokenized residues based on their pairwise distance. The final outcome of this process is a hash table for each binding site. The Tversky index of the pair of hash tables corresponding to the two binding sites is used to quantify the similarity between the binding sites.

### Calculation of sequence similarity

For assessing sequence similarity between antibodies, sequences of the corresponding PDB structures of antibodies present in the Ab-Ligity and SARS-CoV-2 dataset were retrieved from the Protein Data Bank. Full-length variable domain sequence data, as well as CDR sequence data were also extracted from the IEDB (www.iedb.org) (*14*).

#### 1. All-against-all pairwise sequence identity of full-length heavy and light-chain variable domain sequences aligned using MUSCLE

Heavy and light-chain sequences were aligned using MUSCLE (Multiple Sequence Alignment with Log-Expectation) (*16*) which is a general algorithm not restricted to only antibody or T-cell receptor sequences. Sequence identity between each sequence and every other sequence in the alignment was estimated by counting the number of identical residues divided by either the length of the shorter sequence or the length of the longer sequence.

#### 2. All-against-all pairwise sequence identity of full-length heavy and light-chain variable domain sequences aligned using ANARCI

ANARCI (*17*) works by aligning the query sequence to a database of Hidden Markov Models (HMMs). Each HMM is based on the putative germline sequences of a particular variable domain type for a specific species. This is followed by annotating the aligned residues according to one of the numbering schemes (Kabat, Chothia, IMGT, Martin) (*44*). The heavy and light-chain sequences were aligned using ANARCI, and the sequence identity calculated in the same manner as done for MUSCLE-aligned sequences.

#### 3. All-against-all pairwise sequence identity of the CDR loop regions

From the output of ANARCI, the aligned sequences of each CDR loop region were extracted into separate FASTA files. Pairwise sequence identity for each of the six CDR regions was calculated by dividing the number of identical residues by the minimum length of the aligned sequences.

#### 4. All-against-all pairwise CDR k-mer scores

Apart from calculating the pairwise sequence identity of the CDR loops, we also used another approach to estimate the similarity between CDR sequences. This method is an alignment-free k-mer approach currently implemented in the TCRMatch algorithm (*18*). In this approach, each sequence within the pair is broken down into sets of k-mers, with k ranging from 1 to length of the shortest sequence. K-mer pairs between both sequences are compared and a series of calculations ultimately lead to a final ‘k-mer’ score between 0 and 1.

### Calculating structural similarity of CDR loops

Structural similarity was assessed by calculating the root mean square deviation (RMSD) of backbone C_α_ atoms in each CDR loop between pairs of antibodies. CDR loops were defined based on the IMGT numbering scheme. For antibodies in the Ab-Ligity dataset, the RMSD was calculated on the modeled antibody structures rather than on the experimental structures since, in a real-world scenario, structures of many antibodies are not available, and it is more likely that an antibody model would be used instead. Structures of antibodies present in the SARS-CoV-2 dataset were first processed using SCWRL4 to rearrange side-chain orientations and the RMSD was calculated for each CDR loop on the processed antibody structures.

### Building supervised machine-learning models

#### Splitting dataset into training and test dataset

Each of the three datasets (Ab-Ligity, SARS-CoV-2 and IEDB) contain antibody pairs that are either annotated as binding to the same epitope, and assigned to the positive class (‘1’), or annotated as not binding the same epitope and assigned to the negative class (‘0’). We excluded antibody pairs already present in the Ab-Ligity dataset, from the SARS-CoV-2 and IEDB datasets by identifying and removing antibodies having 100% sequence identity in both their heavy-chain and light-chain sequences with antibodies in the Ab-Ligity dataset. We used different train-test strategies for building and assessing the performance of the machine-learning models. First, using Scikit-learn’s train_test split module along with the StratifiedKFold module, the full Ab-Ligity dataset was then shuffled and split into 75% training and 25% test, with a fixed random seed set to 0 to ensure reproducibility. Since the dataset is imbalanced, the random splitting in a stratified manner was done to ensure that the training and test datasets each have the same percentage of positive and negative classes as in the undivided dataset. In addition to the test set of the Ab-Ligity dataset, we also used the external SARS-CoV-2 and IEDB datasets for testing the performance of the models. Second, each of the full Ab-Ligity, SARS-CoV-2 and IEDB datasets were split into sub-datasets of “high-similarity” and “low-similarity” antibody pairs based on a sequence identity threshold of 80% averaged over the VH and VL sequences. The training component of the full Ab-Ligity dataset, the “high-similarity” subset or the “low-similarity” subsets of Ab-Ligity dataset were considered as different training datasets, and the “high-similarity” and “low-similarity” subsets of the SARS-CoV-2 or IEDB datasets were considered as different test datasets. Third, the full SARS-CoV-2 dataset, its “high-similarity” subset, or “low-similarity” subset were used as separate training datasets, while the “high-similarity and “low-similarity” subsets of the Ab-Ligity dataset were used as separate test datasets.

#### Training the binary classifiers

We trained five machine-learning classifiers, Random Forest (RF), Logistic Regression (LR), Gaussian Naive Bayes (GNB), XGBoost (XGB) and a Feed Forward Neural Network (FFNN) in order to build models capable of predicting if a pair of antibodies bind to the same epitope based on selected antibody features. We used the scikit-learn package to implement and train the RF, LR and GNB classifiers. The XGB classifier was implemented using the XGBoost Python package and the FFNN classifier implemented in Tensorflow with Keras as the application programming interface. Prior to training, we carried out normalization of the data present only in the training dataset using scikit-learn’s StandardScaler module. This module calculates the mean and variance of each feature, from the values present only in the training dataset, and then transforms each feature by subtracting the mean and dividing by the variance. Values of features in the test data were normalized by using the scaling values computed on the training dataset.

For the RF classifier, the number of decision trees was set to 100, and the maximum depth of each decision tree was set to 10. To decide the best split, the ‘entropy’ criterion was selected, and the number of features to be considered at each split was set to the square root of the number of features included. A random state of 0 was specified to ensure the reproducibility of results since the results vary due to the bootstrapping of the samples as well as the features considered at each split in the decision tree. GNB and LR models were trained using the default hyperparameters of scikit-learn.

For the XGB and FFNN classifiers, hyperparameter tuning was carried out to identify the best combination of hyperparameters. In order to test combinations of hyperparameters for training the XGB classifier, a grid search with stratified 10-fold cross-validation was performed on the training dataset over the following parameters (i). maximum depth: 3,6. (ii). number of estimators: 100, 150, 200. (iii). Learning rate: 0.05, 0.1,0.2,0.3. The final set of hyperparameters chosen was 200 for number of estimators, maximum depth of 6 and a learning rate of 0.1.

Hyperparameter tuning for the FFNN classifier was carried out in two steps. First, a random search stratified 2-fold cross-validation on the training dataset, which tries out random combinations of user-provided hyperparameters. The following hyperparameters were tested during this step. (i). Optimizer: Stochastic Gradient Descent (SGD), Adam, Adagrad, Momentum. (ii). Number of epochs: 10, 20, 40. (iii). Learning rate: 0.0001, 0.005, 0.001,0.005. (iv). L2 regularization: 0.01, 0.02. (v). Dropout rate: 0.2, 0.5, 0.8. (vi). Number of Neurons: 5, 10,20, 30, 40, 50, 60, 70. (vii). Batch size: 40, 64, 100. (viii). Number of hidden layers: 3, 4, 5. In the second step, a grid search stratified 5-fold cross-validation was performed on the training data over all possible combinations of these hyperparameters: (i). Optimizer: SGD, Adam. (ii). Number of epochs: 20, 30, 50. (iii). Learning rate: 0.005, 0.001, 0.05. (iv). L2 regularization: 0.01. (v). Dropout rate: 0.5, 0.8. Number of Neurons: 20, 30. (vii). Batch size: 64, 128. The best combination of hyperparameters after monitoring the loss on the training set and validation set was found to be with SGD as the optimizer, number of epochs set to 70, number of hidden layers equal to 4, learning rate of 0.001, dropout rate of 0.5, number of neurons set to 30, batch size of 128 and L2 regularization 0.01.

### Developing a command line tool for BCRMatch

Models were trained separately on the Ab-Ligity and IEDB datasets and serialized as pickle files. These models, along with the code, are included with the standalone (https://github.com/IEDB/BCRMatch). Models will be periodically retrained with updated datasets and will be made available via the IEDB Downloads server as well as through updated container images.

## ACKNOWLEDGMENTS

We are grateful to the members of the IEDB, the Bioinformatics Core, and the IT team at the La Jolla Institute for Immunology for their help. We also thank Dr. Wing Ki Wong, the author behind the Ab-Ligity study, for sharing the modelled structures of the antibodies, as well as their predicted paratopes. The research described in this study was supported by the National Institute of Allergy and Infectious Diseases of NIH award no. 75N93019C00001.

## Data and materials availability

All data are available in the main text and supplementary information. The code used to perform the analysis described in this paper, and additional sequence and structural data are available upon request.

