## Supplementary Materials for "Characterizing antibodies binding to the same epitope reveals limited contribution by heavy chain CDR3 sequences relative to other CDRs"

Jarjapu Mahita et al

This PDF file includes:
Supplementary Text
Figs. S1 to S21

**Supplementary Text**

Investigating the impact of the similarity between the Ab-Ligity training and test data on predictive performance

To assess the impact of the similarity of antibody pairs in the Ab-Ligity training dataset to the antibody pairs in the Ab-Ligity test dataset, we defined a similarity of each antibody pair (A,B) in the test dataset to a given antibody pair (C, D) in the training data as:

max_score = max((A-C)+(B-D), (A-D)+(B-C)),

where A-C is the percent sequence identity between antibodies A and C, averaged over the full-length heavy-chain and light-chain variable domain sequences, B-D is the average percent sequence identity between antibodies B and D.

The overall similarity of the antibody pair (A, B) in the test dataset to the training dataset was defined as the maximal similarity across all antibody pairs in the training dataset.

$max\_sim_{A,B} = max[max\_score_1,\ldots,max\_score_i]$

where $max\_score_i$ is the maximal similarity of antibody pair (A,B) to the ith antibody pair in the training dataset.

The above steps were repeated for all antibody pairs in the Ab-Ligity test dataset.

We then created different subsets of the test dataset by excluding antibody pairs above a certain threshold. For example, for one subset, we excluded antibody pairs having a similarity of more than or equal to 99% to the training dataset. For each subset, we evaluated the performance of the machine-learning models through 1000 bootstrap iterations, using different feature combinations, as shown in the figure below (Fig. S_S1). We observed that models trained using sequence features showed lower performance than models trained using a combination of sequence and structural features, with decreasing sequence similarity between the training and test datasets. The mean ROC-AUC value for the LR model trained using only CDR sequence identity features is 0.89 when assessed on the subset of the test dataset having the highest similarity to the training dataset (the '<100 subset', Fig. S_S1). The performance of this LR model decreases to 0.76 when evaluated on the '<90' subset of the test dataset using only CDR sequence features (Fig. S_S1). However, when the LR model is trained using a combination of CDR sequence identity and the paratope shape similarity features represented by the G-LoSA score, the model has a mean ROC-AUC value of 0.92 on the '<100' subset, and 0.81 on the '<90' subset of the test dataset, respectively (Fig. S_S1). These observations reflect the increased contribution of structural features in predictive performance as the sequence similarity between the training and test datasets decreases.

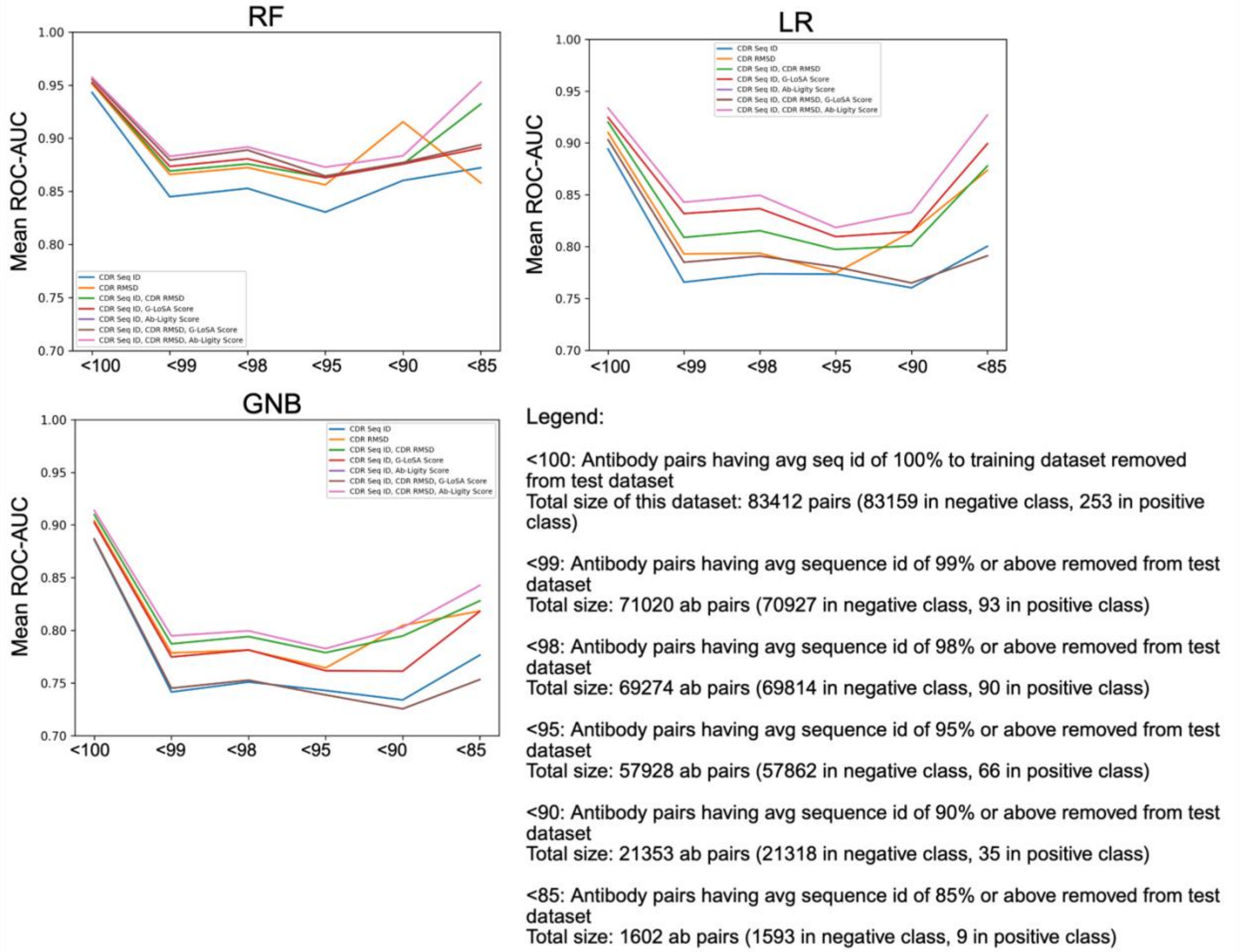


**Fig. S_S1.** Mean ROC-AUC values of the Random Forest (RF), Logistic Regression (LR), and the Gaussian Naïve Bayes (GNB) models with decreasing sequence similarity between the Ab-Ligity training and test datasets, for various feature combinations.

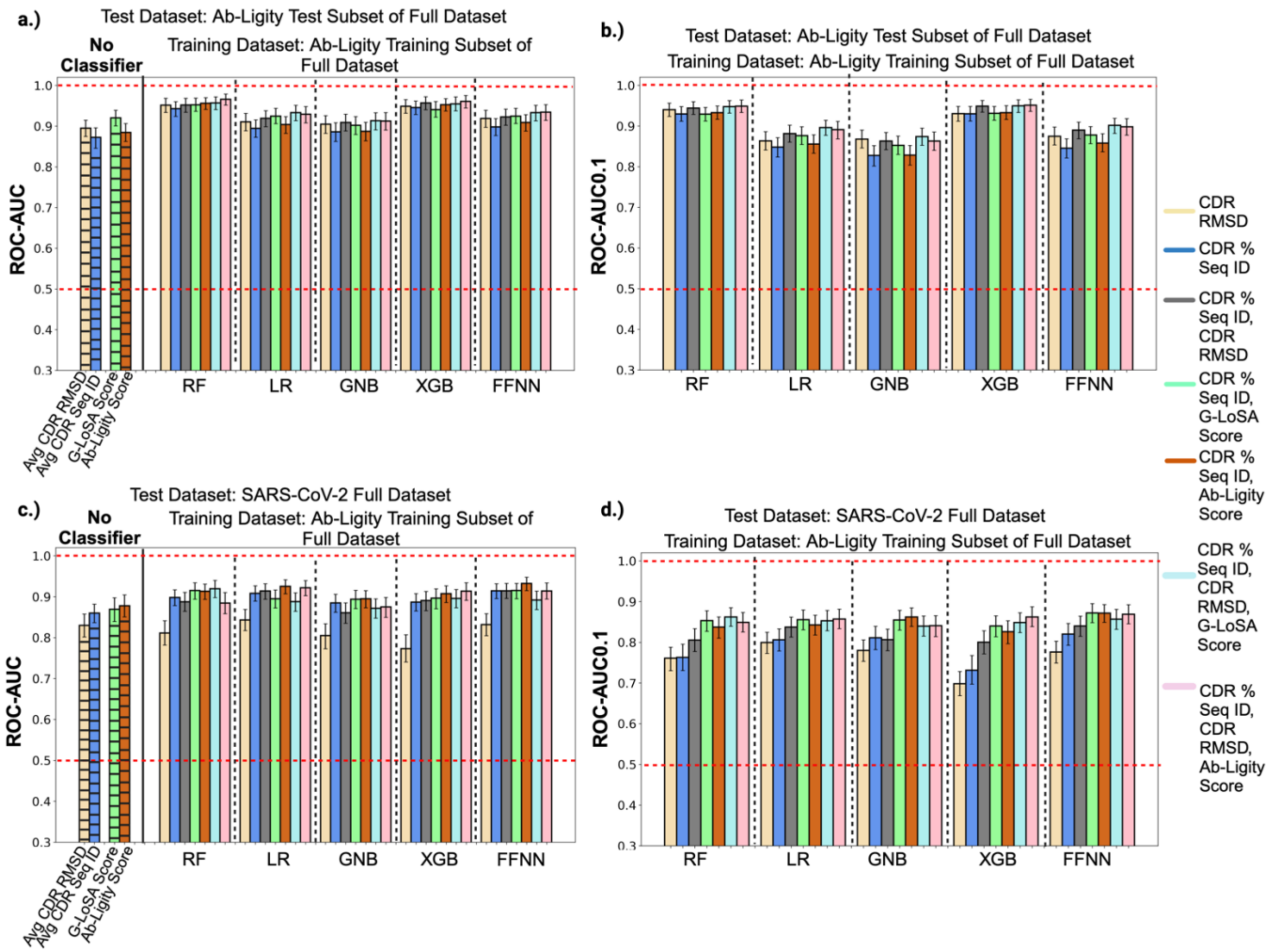


**Fig. S1**. ROC-AUC and ROC-AUC0.1 values of the Random Forest (RF), Logistic Regression (LR), Gaussian Naive Bayes (GNB), XGBoost (XGB) and the Feed Forward Neural Network (FFNN) models when using a combination of sequence and structure features to predict antibodies targeting a common epitope in the Ab-Ligity test dataset (a, b), and the SARS-CoV-2 dataset (c, d). Striped bars represent the mean AUC values when using average CDR RMSD or CDR sequence identity of the six CDR loops, the G-LoSA score, and Ab-Ligity score directly as predictors. Error bars denote the 90% confidence interval (CI).

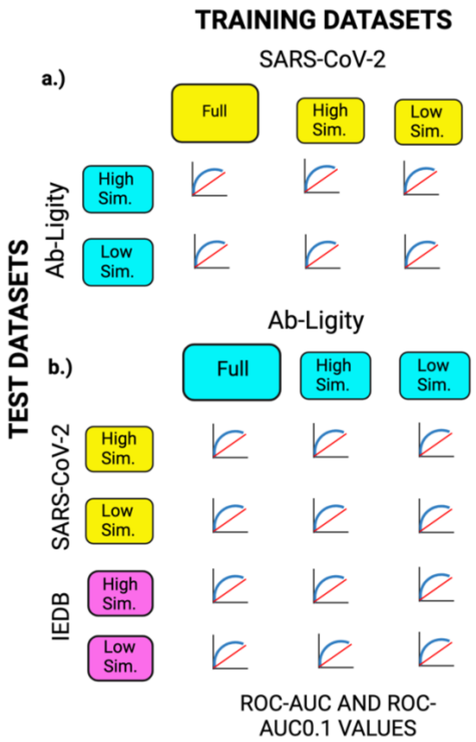


**Fig. S2**. Schematic of the high and low similarity datasets used for training and testing the machine learning models.

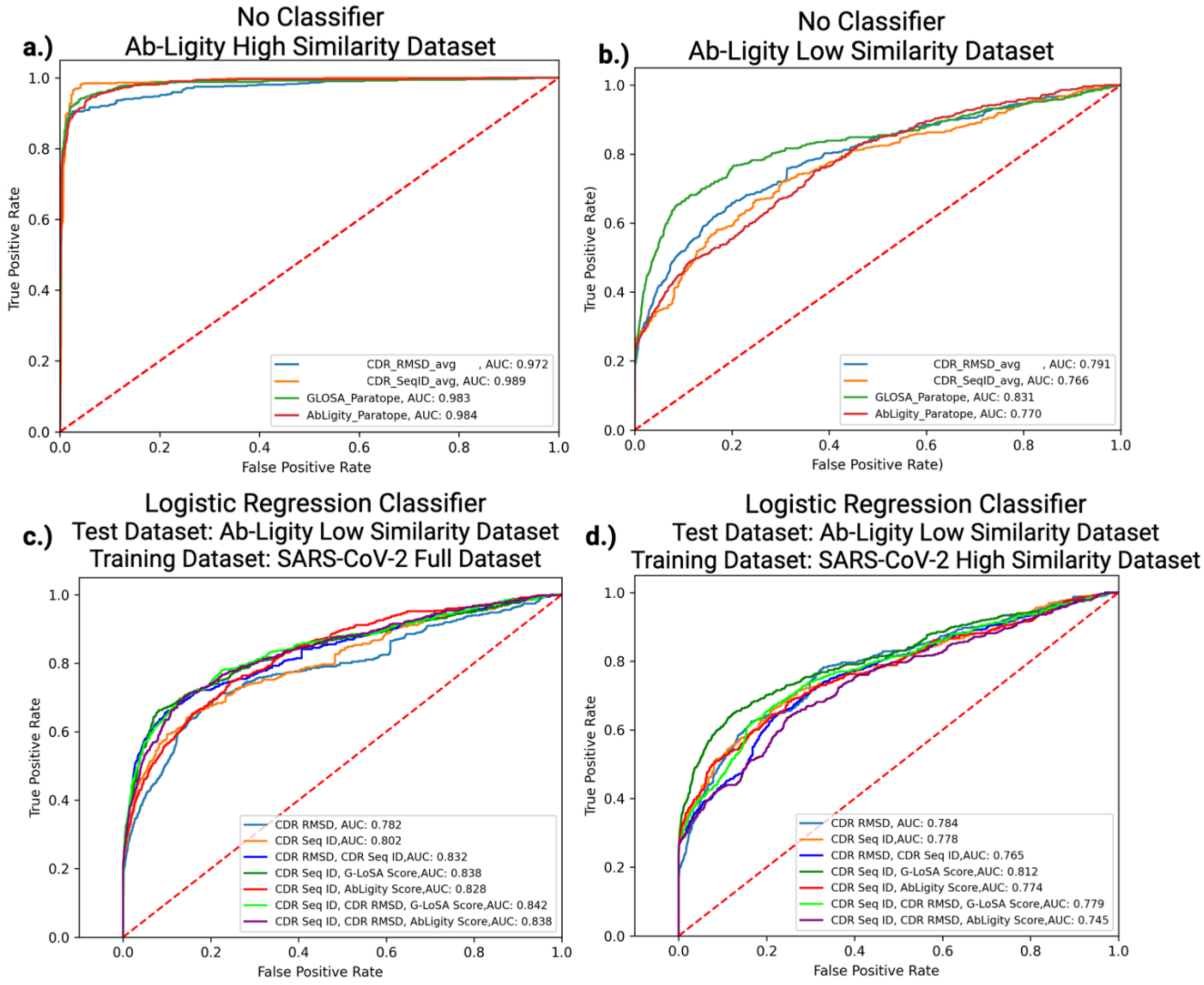
a.)
No Classifier
Ab-Ligity High Similarity Dataset
True Positive Rate
False Positive Rate
CDR_RMSD_avg , AUC: 0.972
CDR_SeqID_avg, AUC: 0.989
GLOSA_Paratope, AUC: 0.983
AbLigity_Paratope, AUC: 0.984
b.)
No Classifier
Ab-Ligity Low Similarity Dataset
True Positive Rate)
False Positive Rate)
CDR_RMSD_avg , AUC: 0.791
CDR_SeqID_avg, AUC: 0.766
GLOSA_Paratope, AUC: 0.831
AbLigity_Paratope, AUC: 0.770
c.)
Logistic Regression Classifier
Test Dataset: Ab-Ligity Low Similarity Dataset
Training Dataset: SARS-CoV-2 Full Dataset
True Positive Rate
False Positive Rate
CDR RMSD, AUC: 0.782
CDR Seq ID,AUC: 0.802
CDR RMSD, CDR Seq ID,AUC: 0.832
CDR Seq ID, G-LoSA Score,AUC: 0.838
CDR Seq ID, AbLigity Score,AUC: 0.828
CDR Seq ID, CDR RMSD, G-LoSA Score,AUC: 0.842
CDR Seq ID, CDR RMSD, AbLigity Score,AUC: 0.838
d.)
Logistic Regression Classifier
Test Dataset: Ab-Ligity Low Similarity Dataset
Training Dataset: SARS-CoV-2 High Similarity Dataset
True Positive Rate
False Positive Rate
CDR RMSD, AUC: 0.784
CDR Seq ID,AUC: 0.778
CDR RMSD, CDR Seq ID,AUC: 0.765
CDR Seq ID, G-LoSA Score,AUC: 0.812
CDR Seq ID, AbLigity Score,AUC: 0.774
CDR Seq ID, CDR RMSD, G-LoSA Score,AUC: 0.779
CDR Seq ID, CDR RMSD, AbLigity Score,AUC: 0.745

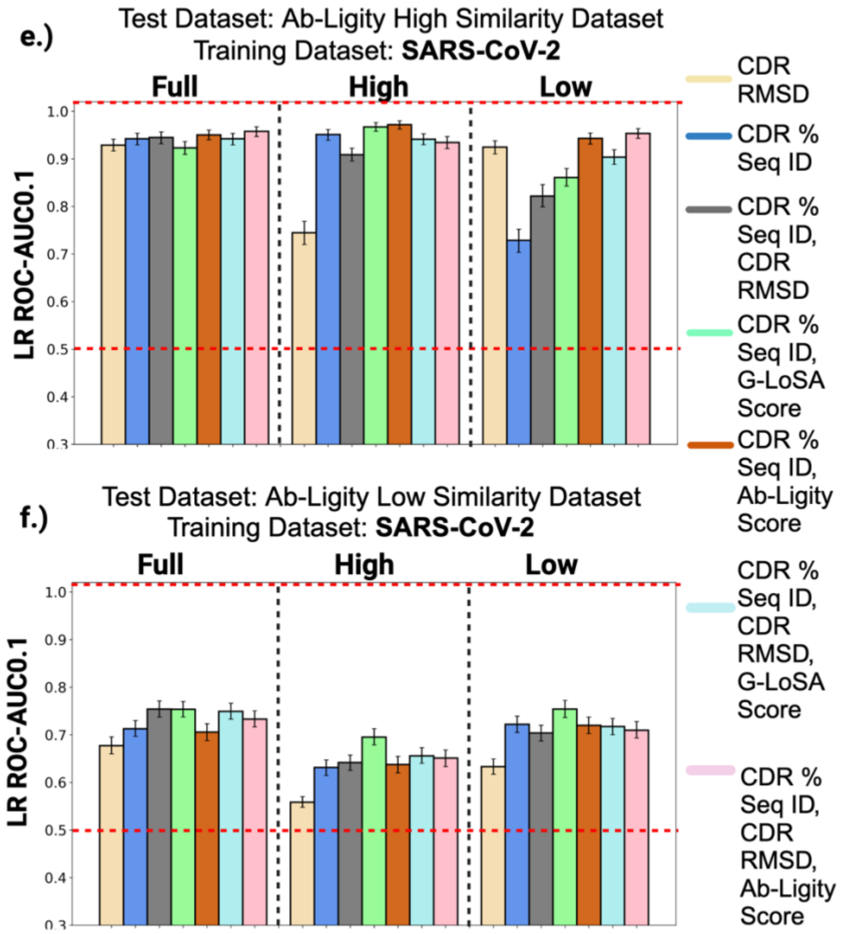


**Fig. S3**. ROC plots showing the performance of different predictors on the Ab-Ligity high similarity dataset (a.) and the low similarity dataset (b). ROC curves of the Logistic Regression models tested on different combinations of features in the Ab-Ligity low similarity dataset after training on the full SARS-CoV-2 dataset (c.) or the SARS-CoV-2 high similarity dataset. e,f.) ROC-AUC0.1 values of the LR models tested on the Ab-Ligity high and low similarity datasets.

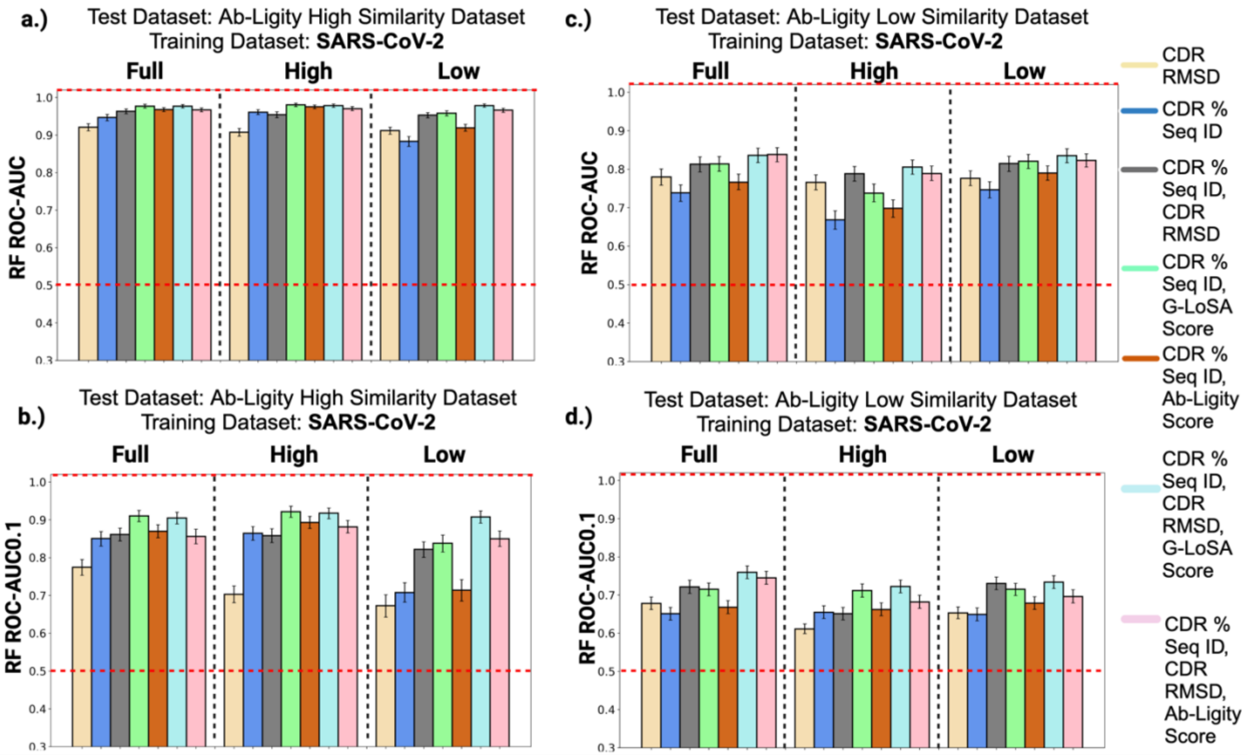


**Fig. S4**: a, b.)ROC-AUC and ROC-AUC0.1 values of the Random Forest (RF) models tested on the Ab-Ligity high similarity dataset after being trained on either the SARS-CoV-2 full, high or low similarity datasets. c,d.) RF performances on the Ab-Ligity low similarity dataset.

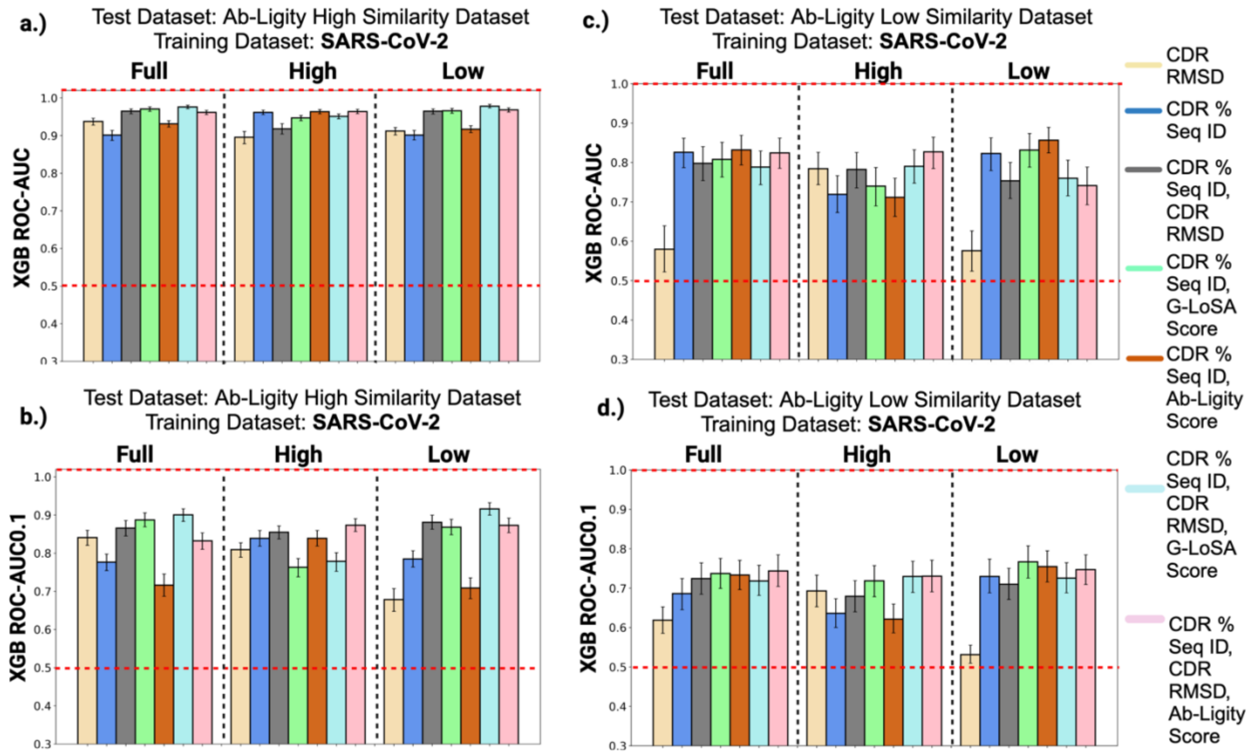


**Fig. S5**. a, b.)ROC-AUC and ROC-AUC0.1 values of the XGBoost (XGB) models tested on the Ab-Ligity high similarity dataset after being trained on either the SARS-CoV-2 full, high or low similarity datasets. c,d.) XGB model performances on the Ab-Ligity low similarity dataset.

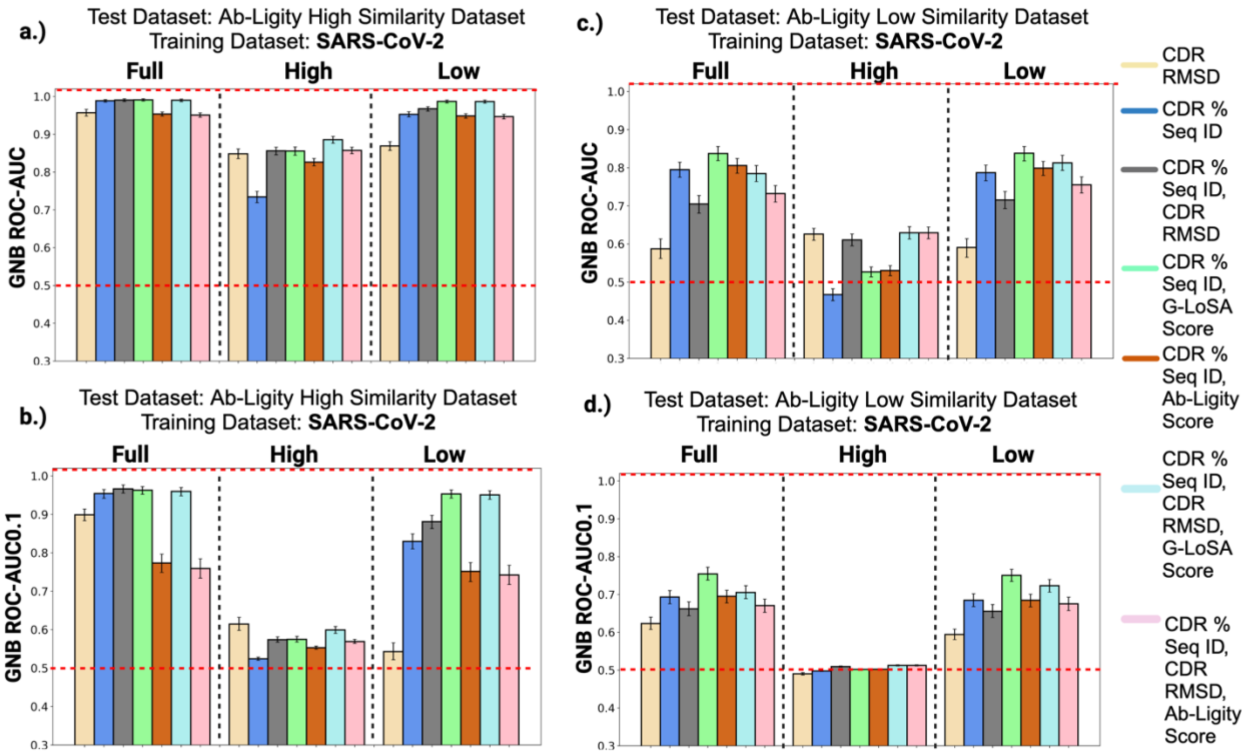


**Fig. S6**. a, b.)ROC-AUC and ROC-AUC0.1 values of the Gaussian Naive Bayes (GNB) models tested on the Ab-Ligity high similarity dataset after being trained on either the SARS-CoV-2 full, high or low similarity datasets. c,d.) GNB performances on the Ab-Ligity low similarity dataset.

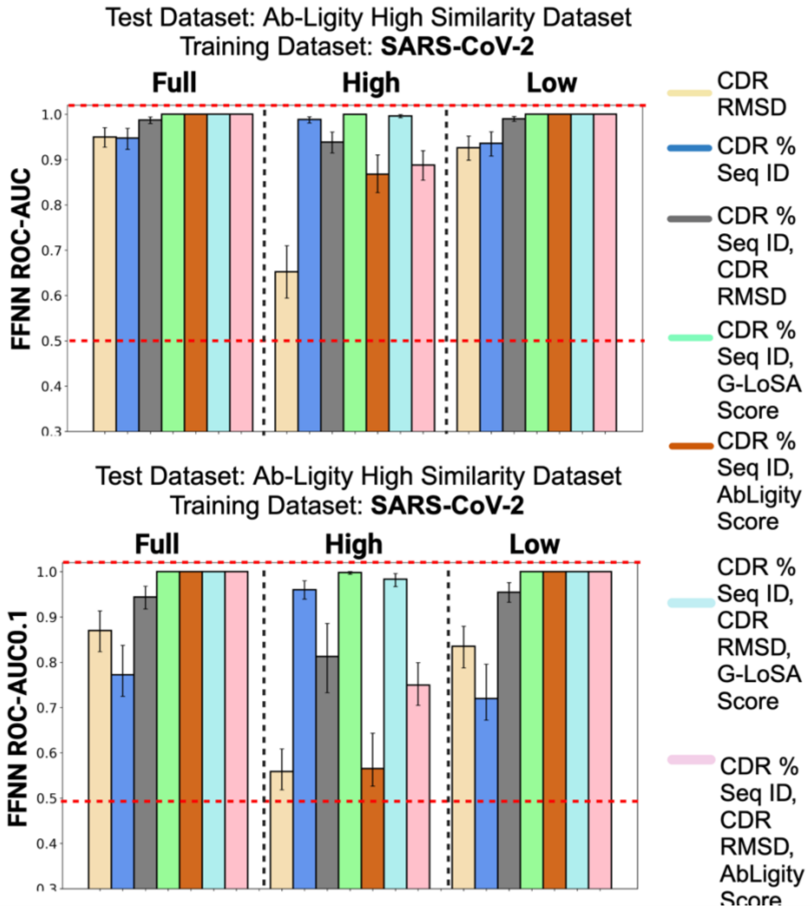


**Fig. S7**. ROC-AUC and ROC-AUC0.1 values of the Feed Forward Neural Network (FFNN) models tested on the Ab-Ligity high similarity dataset after being trained on either the SARS-CoV-2 full, high or low similarity datasets.

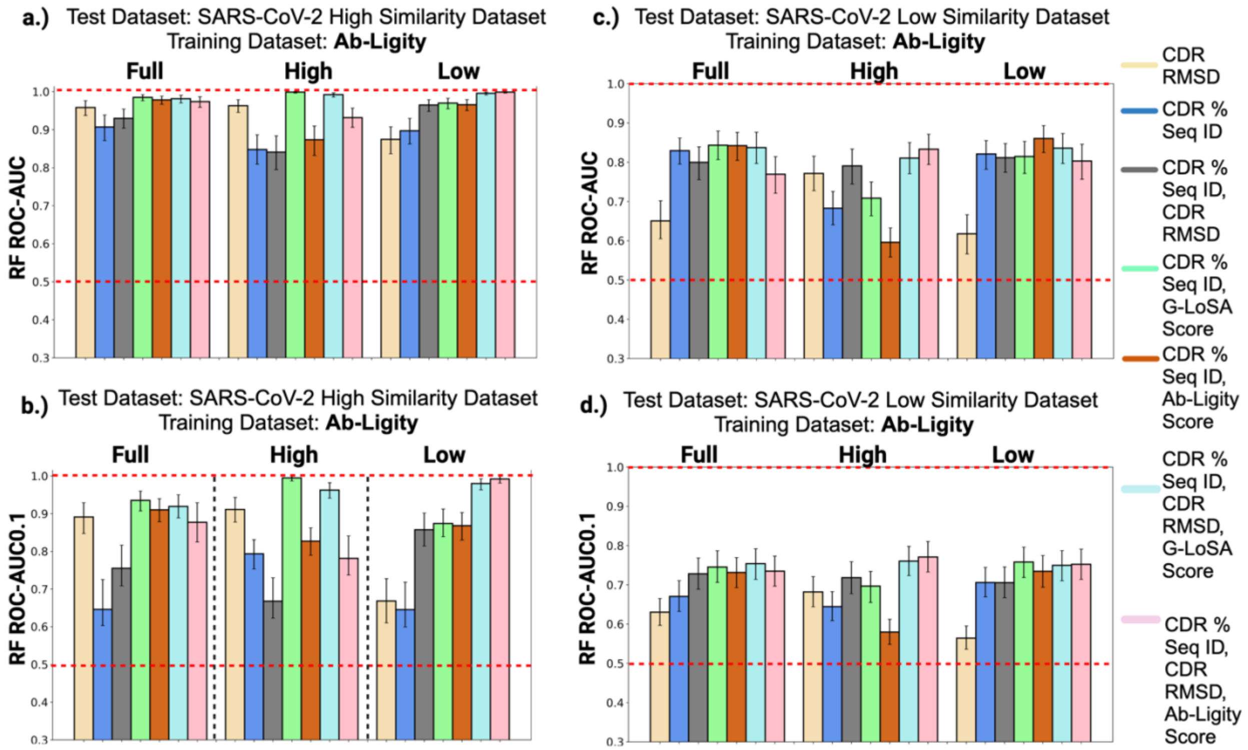


**Fig. S8**. a, b.)ROC-AUC and ROC-AUC0.1 values of the Random Forest (RF) models tested on the SARS-CoV-2 high similarity dataset after being trained on either the Ab-Ligity full, high or low similarity datasets. c,d.) RF performances on the SARS-CoV-2 low similarity dataset.

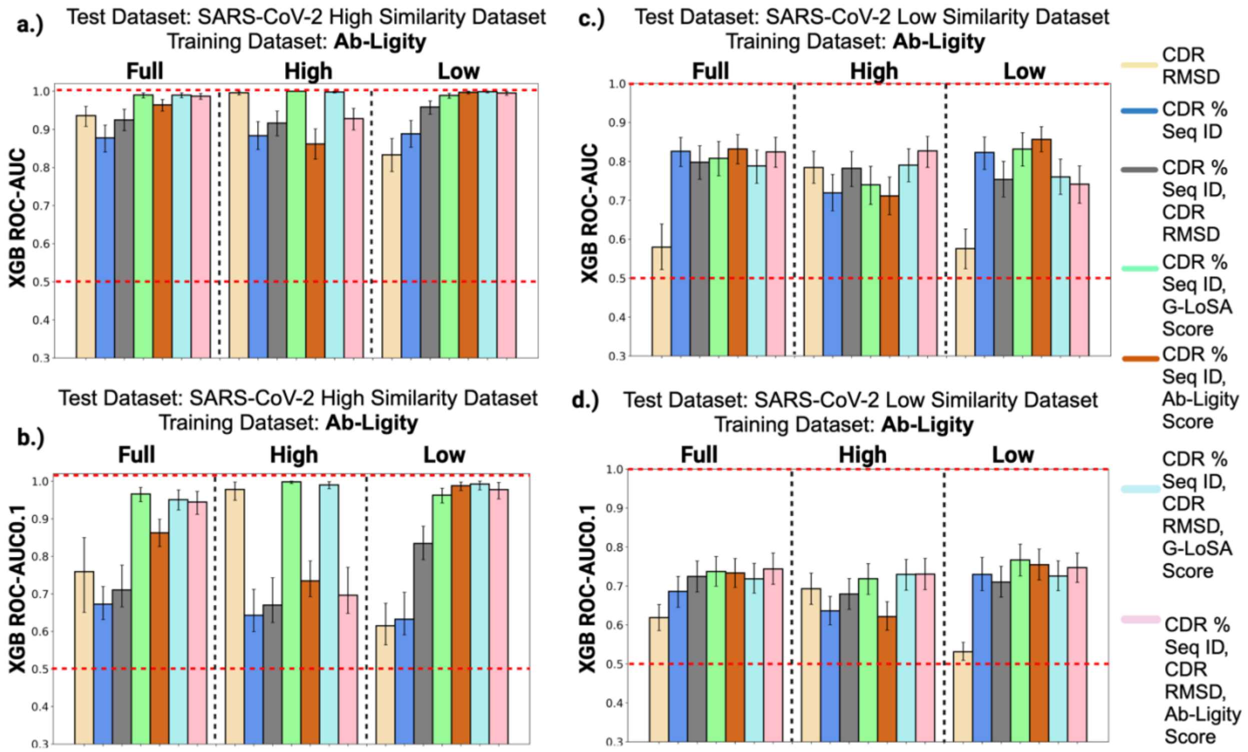


**Fig. S9**. a, b.)ROC-AUC and ROC-AUC0.1 values of the XGBoost (XGB) models tested on the SARS-CoV-2 high similarity dataset after being trained on either the Ab-Ligity full, high or low similarity datasets. c,d.) Performances of XGB models on the SARS-CoV-2 low similarity dataset.

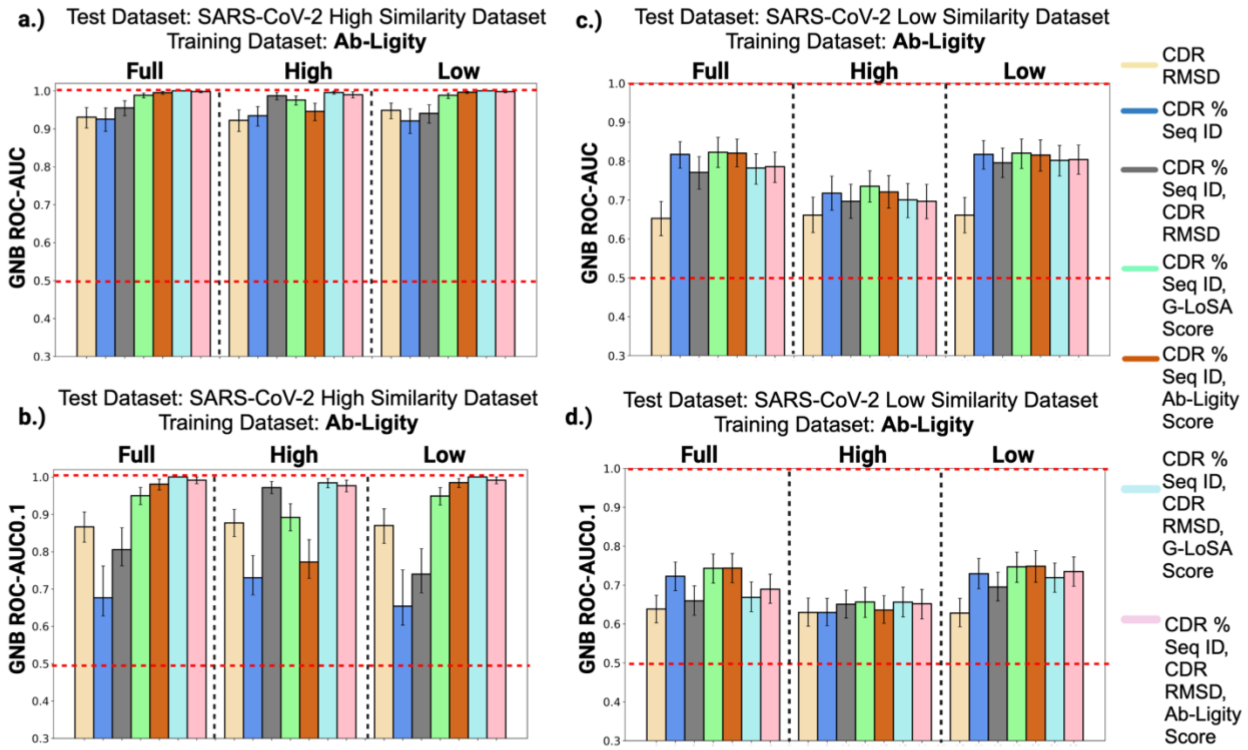


**Fig. S10**. a, b.)ROC-AUC and ROC-AUC0.1 values of the Gaussian Naive Bayes (GNB) models tested on the SARS-CoV-2 high similarity dataset after being trained on either the Ab-Ligity full, high or low similarity datasets. c,d.) Performances of GNB models  on the SARS-CoV-2 low similarity dataset.

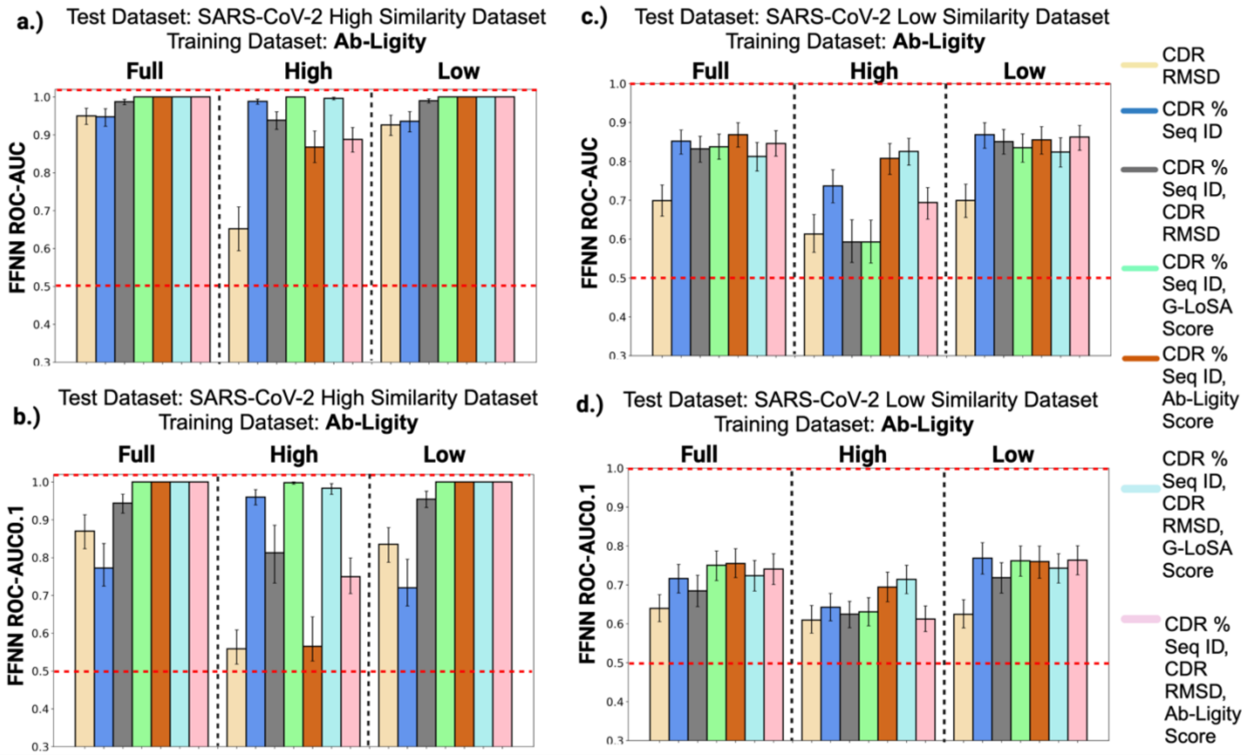


**Fig. S11**. a, b.)ROC-AUC and ROC-AUC0.1 values of the Feed Forward Neural Network (FFNN) models tested on the SARS-CoV-2 high similarity dataset after being trained on either the Ab-Ligity full, high or low similarity datasets. c,d.) Performances of FFNN models on the SARS-CoV-2 low similarity dataset.

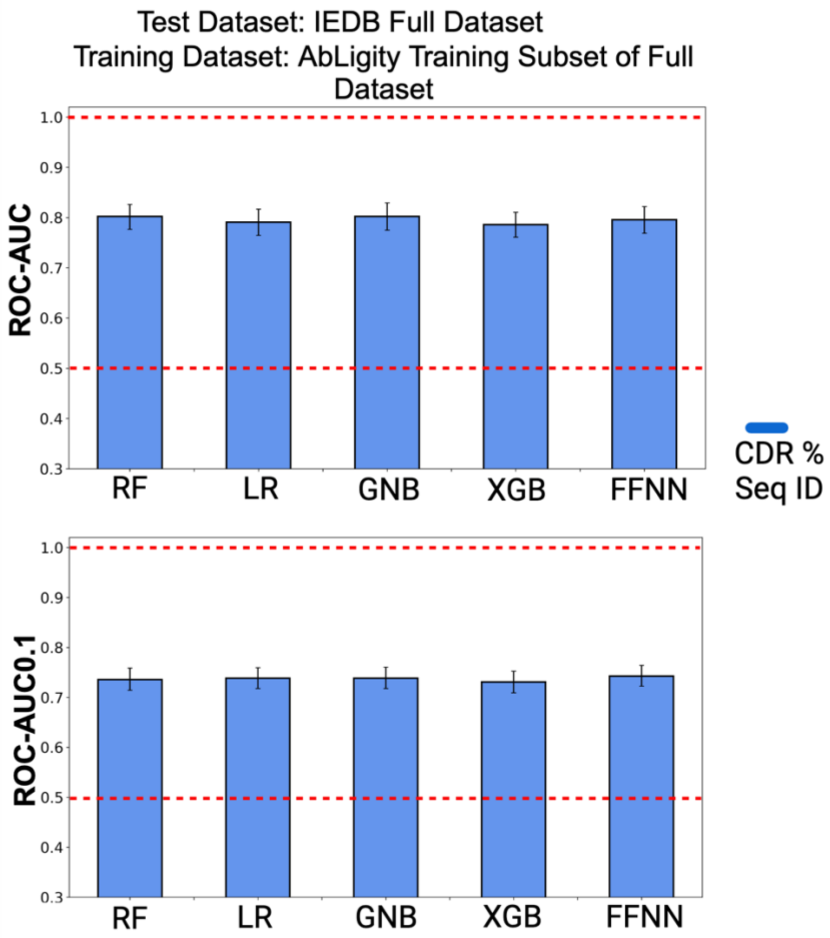


**Fig. S12**. ROC-AUC and ROC-AUC0.1 values of the five ML models tested on the IEDB full dataset using only the CDR sequence identity of the six CDR loops as features.

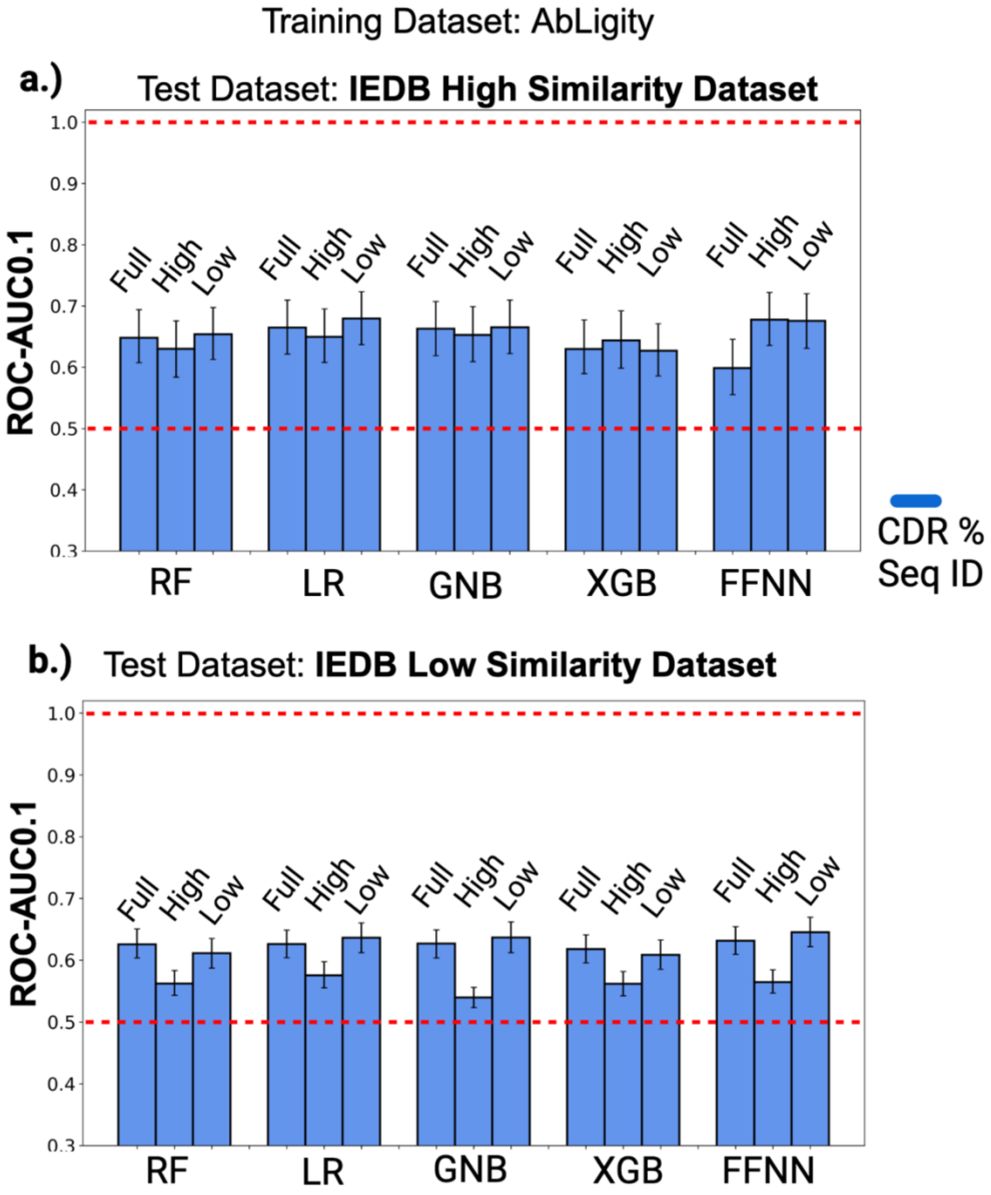


**Fig. S13**. a.) Partial ROC-AUC (ROC-AUC0.1) values of the five machine learning models tested on the IEDB high similarity dataset using the full, high or low similarity Ab-Ligity datasets for training. b.) ROC-AUC0.1 values when the IEDB low similarity dataset is used for testing.

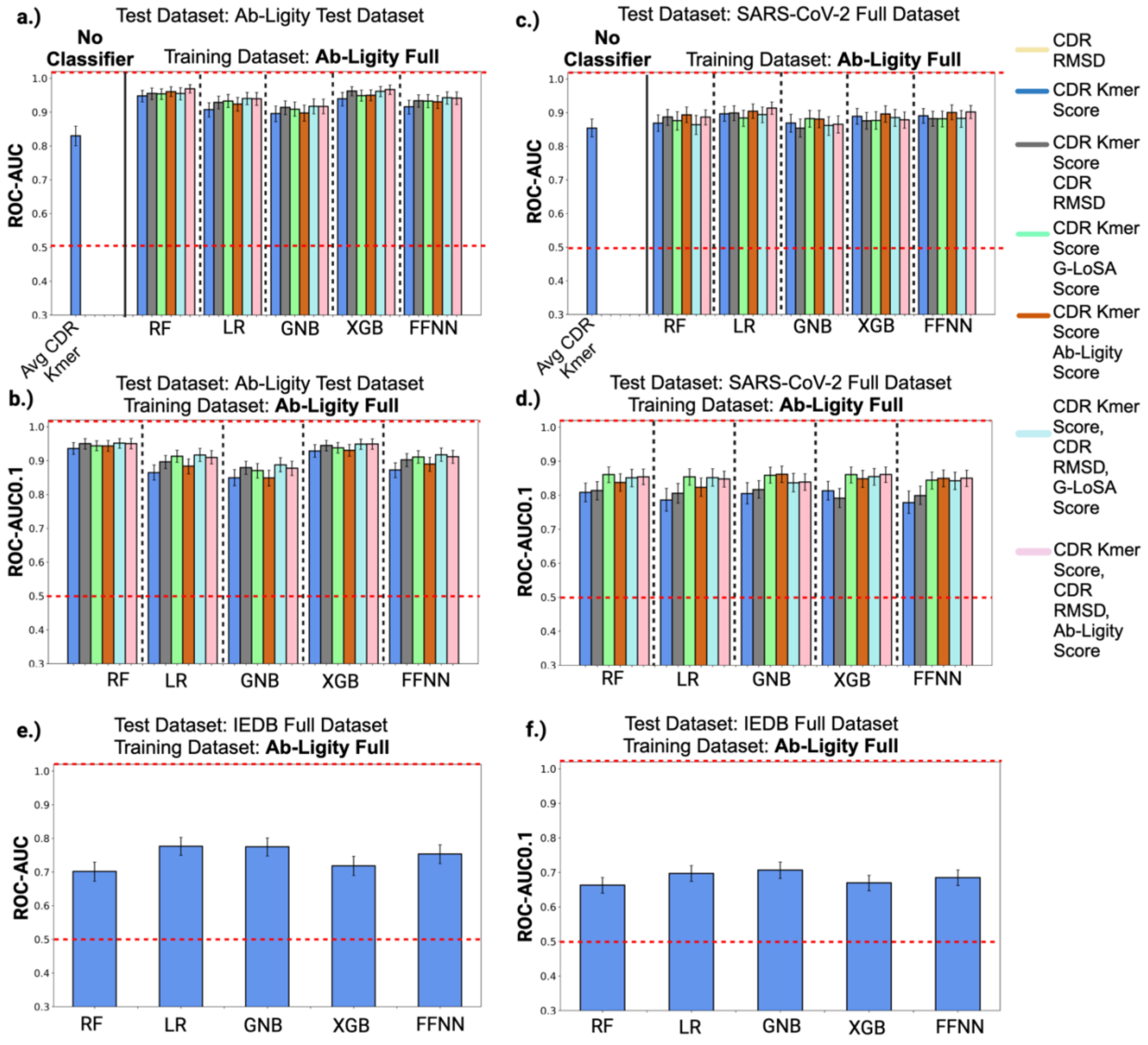


**Fig. S14**. ROC-AUC and ROC-AUC0.1 values of the Random Forest (RF), Logistic Regression (LR), Gaussian Naive Bayes (GNB), XGBoost (XGB) and the Feed Forward Neural Network (FFNN) models when using a combination of sequence and structure features to predict antibodies targeting a common epitope after replacing the CDR sequence identity feature by the CDR k-mer score feature in the Ab-Ligity test dataset (a, b), and the SARS-CoV-2 dataset (c, d). Bars labeled ‘Avg CDR Kmer’ in a) and c) represent the mean AUC value when using average CDR K-mer score of the six CDR loops directly as a predictor. (e, f). ROC-AUC and ROC-AUC0.1 values when models are tested on the IEDB full dataset using only the CDR K-mer scores as features. Error bars denote the 90% confidence interval (CI).

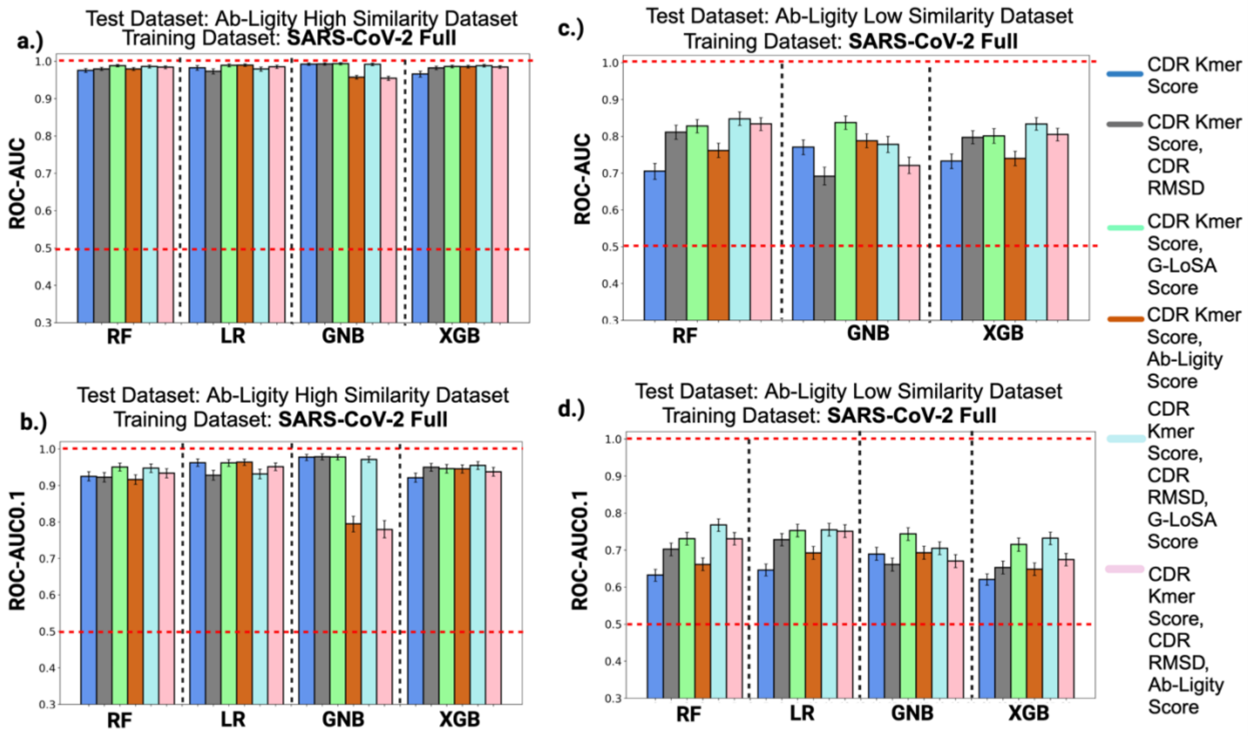


**Fig. S15**. ROC-AUC and ROC-AUC0.1 values of the five ML models tested on the Ab-Ligity high similarity dataset using the full SARS-CoV-2 dataset for training (a,b). c,d).AUC and AUC0.1 values of the models evaluated on the Ab-Ligity low similarity dataset. Note that the CDR sequence identity feature is replaced by the CDR k-mer score feature in these predictions.

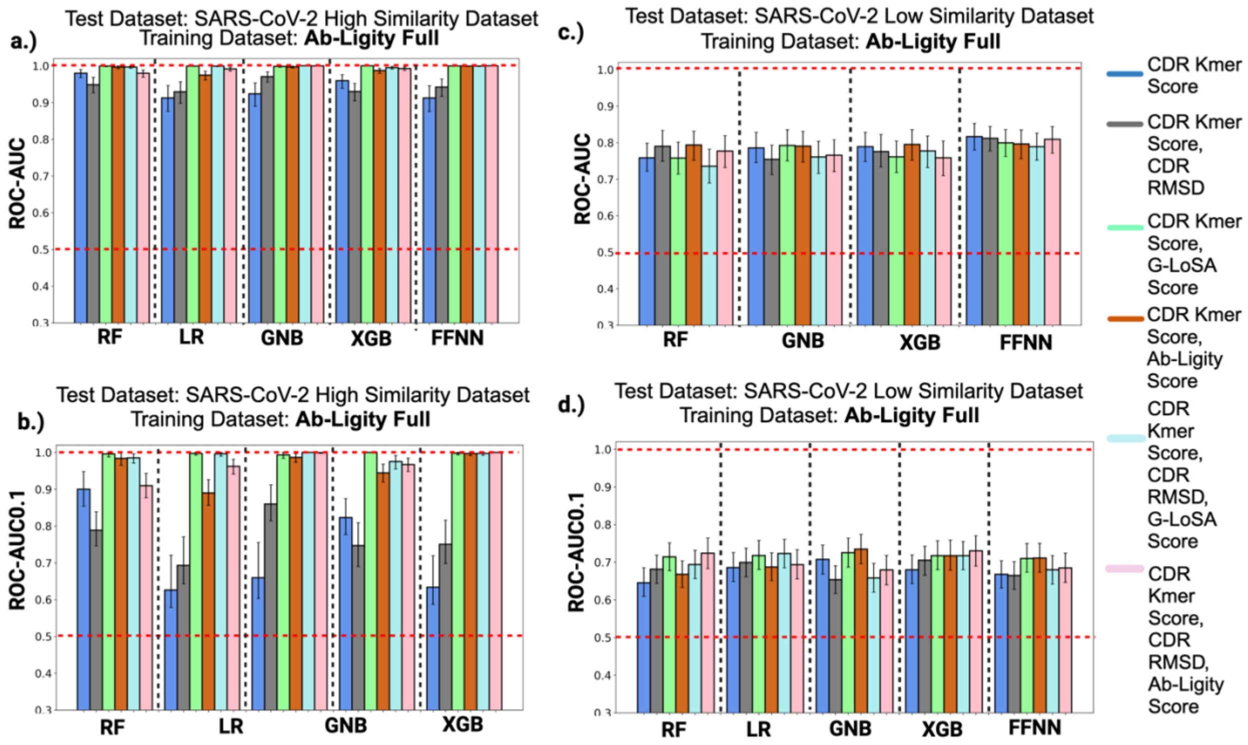


**Fig. S16**. ROC-AUC and ROC-AUC0.1 values of the five ML models tested on the SARS-CoV-2 high similarity dataset using the Ab-Ligity Full dataset for training after substituting the features of CDR sequence identity by the corresponding CDR k-mer score(a, b). c, d.) AUC and AUC0.1 values of the models evaluated on the SARS-CoV-2 low similarity dataset.

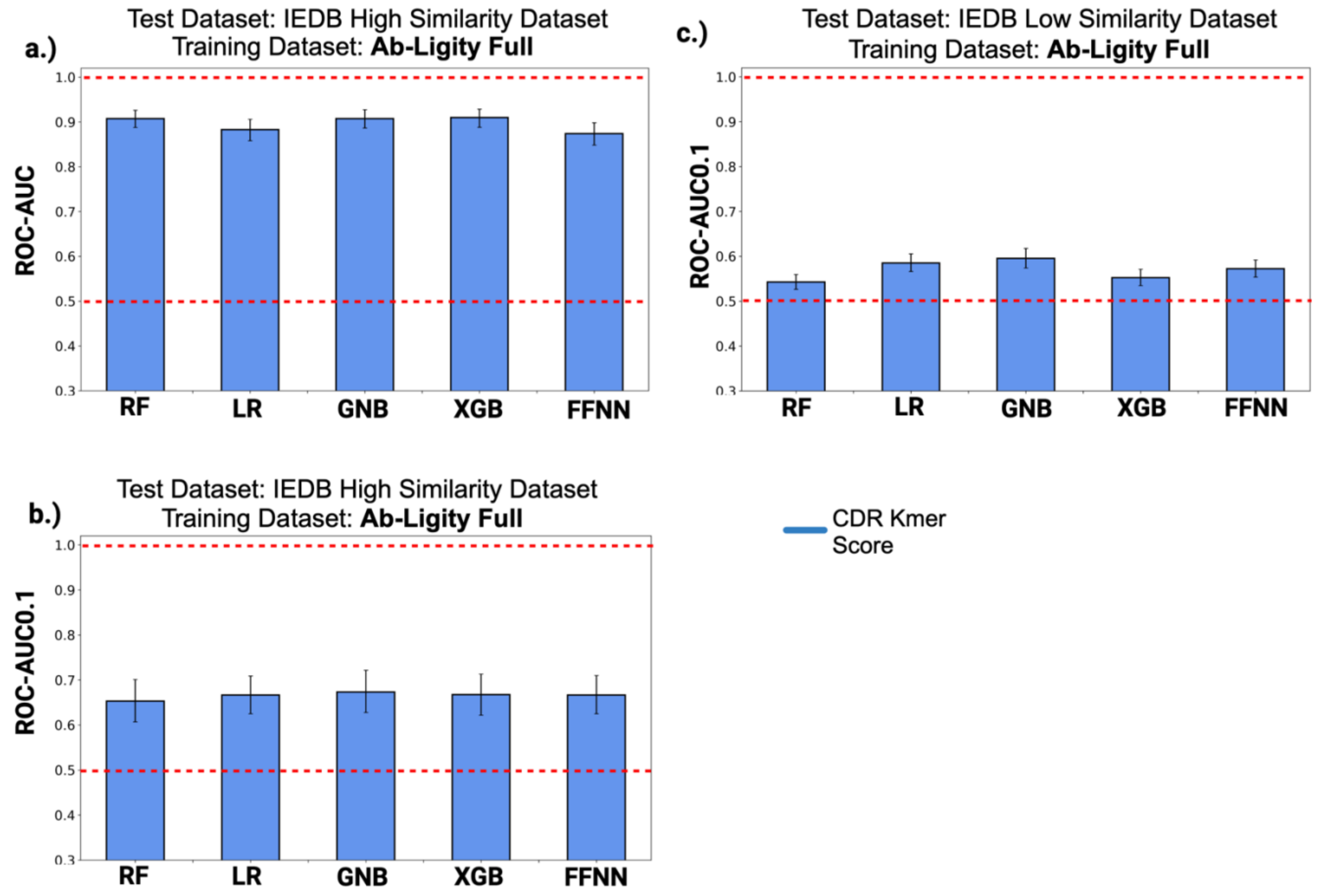


**Fig. S17**. a, b.): ROC-AUC and AUC0.1 values of the five ML models trained on the Ab-Ligity Full dataset and tested on the IEDB high similarity dataset using only the CDR k-mer scores of the six CDR loops as features. c.) AUC0.1 values of the models on the IEDB low similarity dataset.

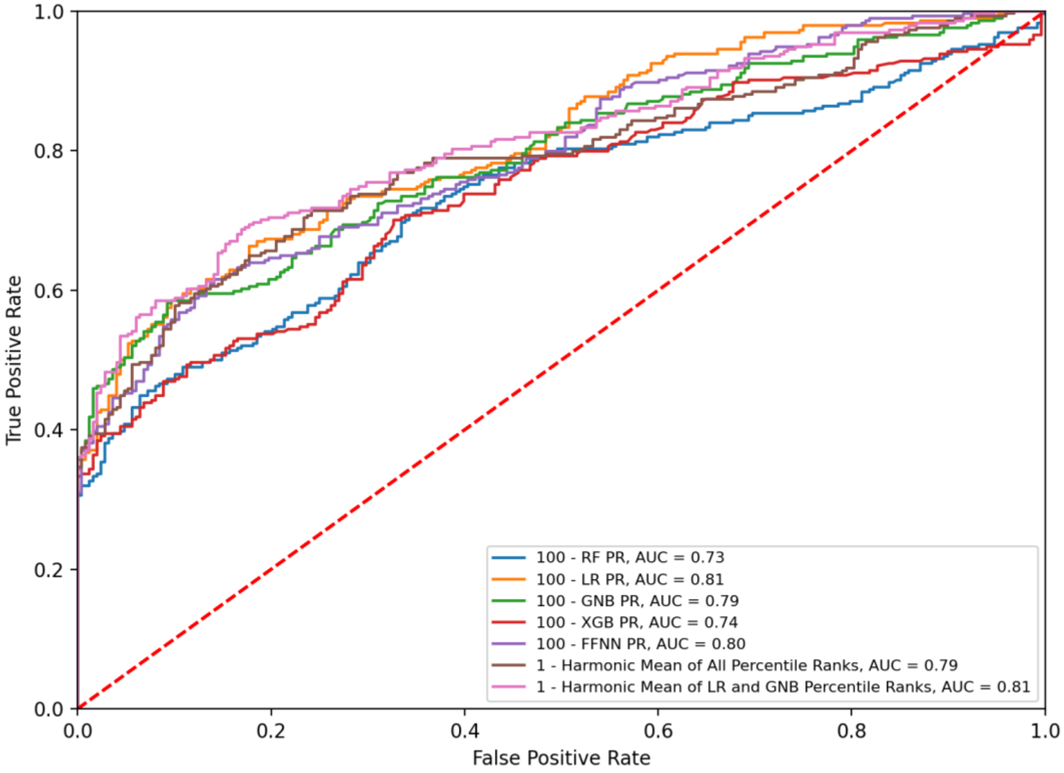


**Fig. S18**: Comparison of ROC-AUC values of the individual percentile ranks of the five ML models versus the harmonic mean of percentile rank of either all five models or only the LR and GNB models, tested on a random subset of the IEDB dataset. PR: Percentile rank.

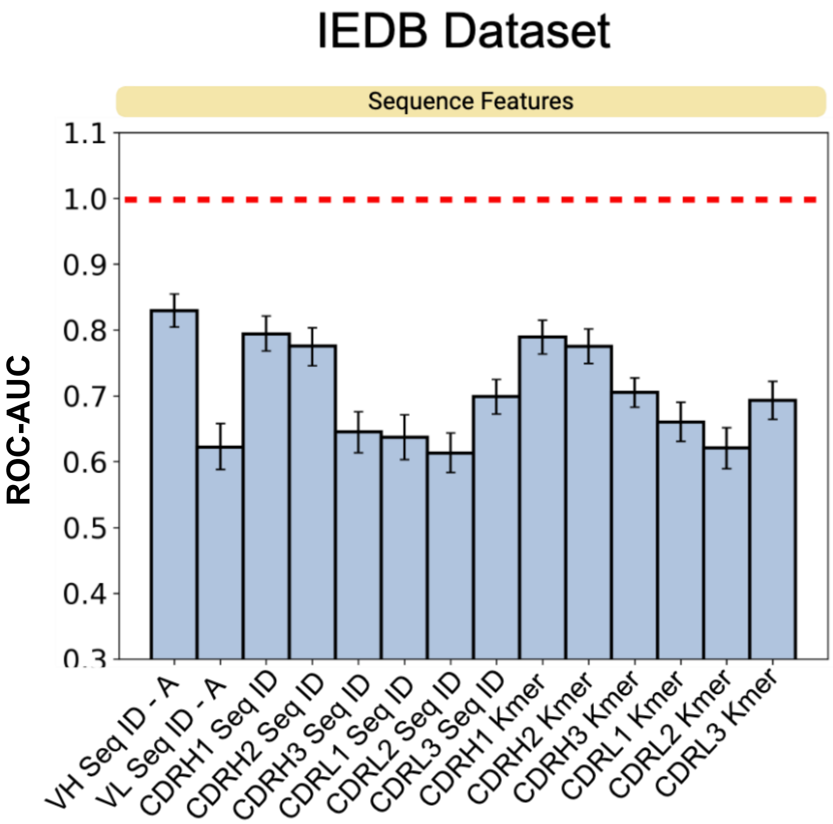


**Fig. S19:** ROC-AUC values when using individual features to predict antibodies targeting a common epitope in the IEDB dataset.

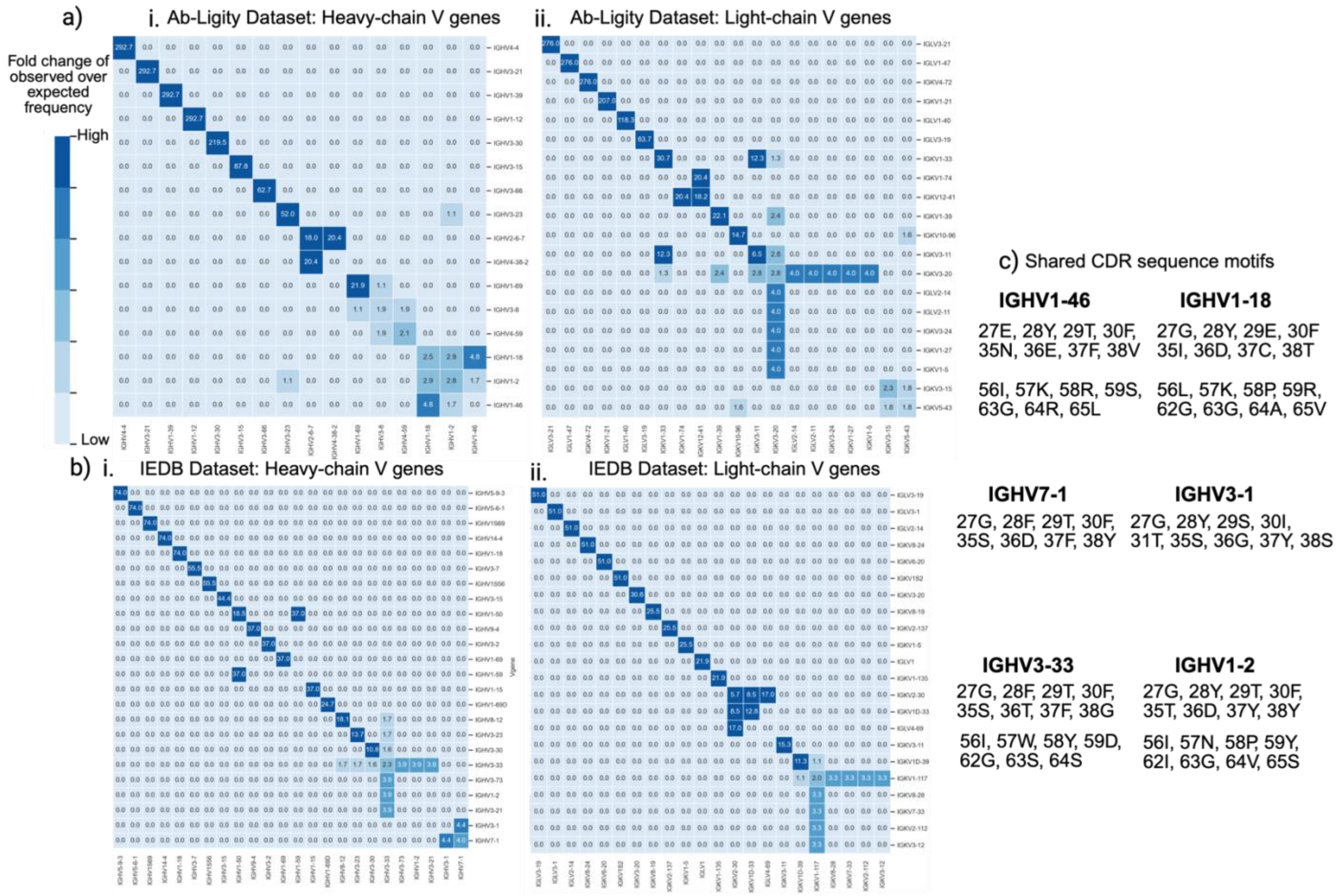


**Fig. S20:** a.) Fold change of observed over expected frequencies of i. heavy-chain V gene pairs, ii. light-chain V gene pairs in the positive class of the Ab-Ligity dataset. b.) Fold change of observed over expected frequencies of i. heavy-chain V gene pairs, ii. light-chain V gene pairs in the positive class of the IEDB dataset. c.) Examples of shared CDR sequence motifs identified in the Ab-Ligity and IEDB datasets. Numbering is as per IMGT scheme.

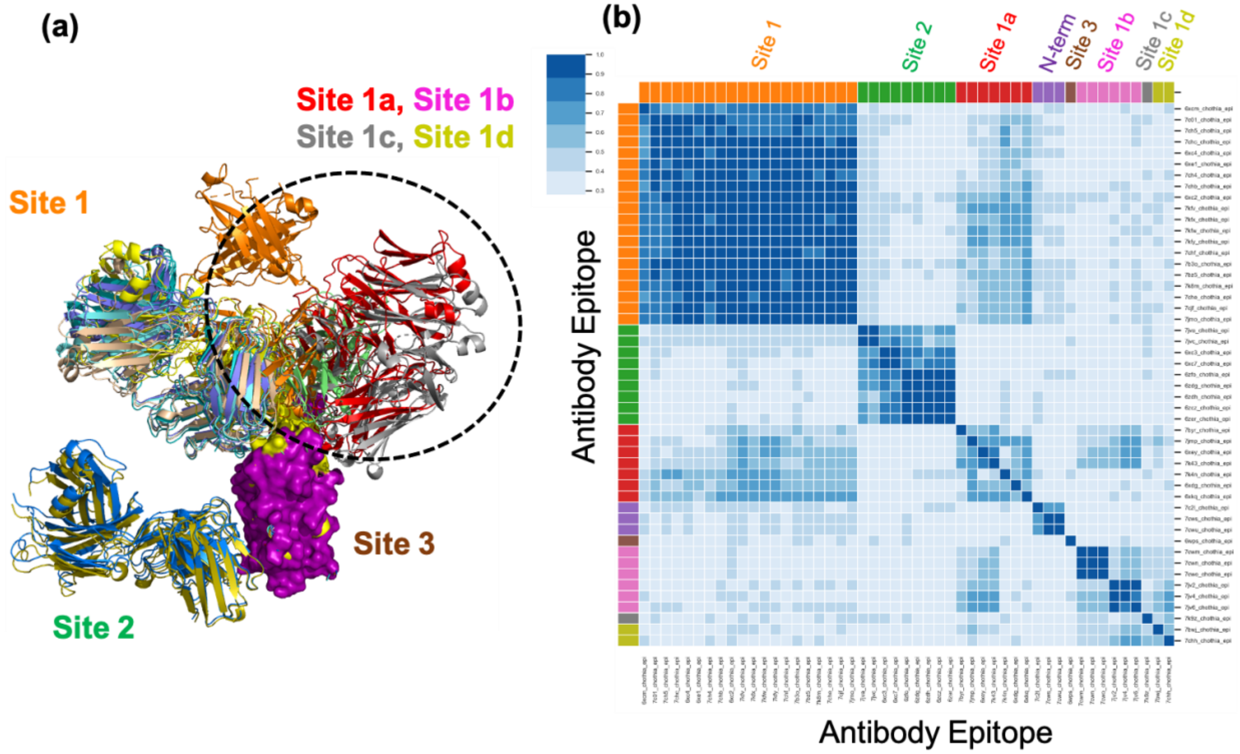


**Fig. S21**: **(a)** Location of different SARS-CoV-2 antibody epitopes on the receptor-binding domain (RBD) of the Spike protein. Epitopes on the N-terminal domain of SARS-CoV-2 are not shown. (**b**) All-against-all comparison of the SARS-CoV-2 antibody epitopes was carried out using the binding site prediction algorithm and the values plotted as a heatmap, highlighting the similarity of overlapping epitopes. Darker the color, higher is the similarity score.
